# Humoral immune response to adenovirus induce tolerogenic bystander dendritic cells that promote generation of regulatory T cells

**DOI:** 10.1101/334508

**Authors:** Thi Thu Phuong Tran, Karsten Eichholz, Patrizia Amelio, Crystal Moyer, Glen R Nemerow, Matthieu Perreau, Franck JD Mennechet, Eric J Kremer

## Abstract

Following repeated encounters with adenoviruses most of us develop robust humoral and cellular immune responses that are thought to act together to combat ongoing and subsequent infections. Yet in spite of robust immune responses, adenoviruses establish subclinical persistent infections that can last for decades. While adenovirus persistence pose minimal risk in B-cell compromised individuals, if T-cell immunity is severely compromised, reactivation of latent adenoviruses can be life threatening. This dichotomy led us to ask how anti-adenovirus antibodies influence adenovirus-specific T-cell immunity. Using primary human blood cells, transcriptome and secretome profiling, and pharmacological, biochemical, genetic, molecular, and cell biological approaches, we initially found that healthy adults harbor adenovirus-specific regulatory T cells (T_regs_). As peripherally induced T_regs_ are generated by tolerogenic dendritic cells (DCs), we then addressed how tolerogenic DCs could be created. Here, we demonstrate that DCs that take up immunoglobulin-complexed (IC)-adenoviruses create an environment that causes bystander DCs to become tolerogenic. These adenovirus antigen-loaded tolerogenic DCs can drive naïve T cells to mature into adenovirus-specific T_regs_. Our results may provide ways to improve antiviral therapy and/or pre-screening high-risk individuals undergoing immunosuppression.

**Author summary:** While numerous studies have addressed the cellular and humoral response to primary virus encounters, relatively little is known about the interplay between persistent infections, neutralizing antibodies, antigen-presenting cells, and the T-cell response. Our studies suggests that if adenovirus–antibody complexes are taken up by professional antigen-presenting cells (dendritic cells), the DCs generate an environment that causes bystander dendritic cells to become tolerogenic. These tolerogenic dendritic cells favors the creation of adenovirus-specific regulatory T cells. While this pathway likely favors pathogen survival, there may be advantages for the host also.

## Introduction

Human adenoviruses (HAdVs), of which there are currently >80 types, typically cause self-limiting respiratory, ocular, and gastro-intestinal tract infections in immunocompetent individuals. After repeated encounters, most young adults generally harbor cross-reactive, long-lived humoral and T-cell responses [1–3] that are thought to work together to efficiently blunt subsequent HAdV-induced morbidity. However, in spite of the robust anti-HAdV immune responses, HAdVs routinely establish decades-long, subclinical infections that are characterized by low level shedding of progeny virions [4,5]. While potential molecular mechanisms by which HAdVs evade the immune response have been proposed [6], we suspected that complementary mechanisms also exist. Of note, in T-cell compromised patients the loss of cellular control of persistent HAdV infection can lead to fulminant and fatal disease [4,5]. It is noteworthy that serological evidence that the patient has been infected by a given HAdV type before hematopoietic stem cell transplantation is predictive of escape from the same HAdV type during immune suppression [7].

While T-cell therapy has shown a notable potential to prevent HAdV disease in immunocompromised patients [8,9], immunoglobulin therapy has had remarkably little impact [4]. Due to omnipresent anti-HAdV antibodies, it is not surprising that immunoglobulin-complexed HAdVs (IC-HAdVs) are detected in some patients with HAdV disease [10–12]. In a broader view, immunoglobulin-complexed viruses can form during prolonged viremia, secondary infections, primary infections when a cross-reactive humoral response exists, and in the presence of antibody-based antiviral therapy. It is important to note that IC-HAdVs are potent stimulators of human dendritic cell (DC) maturation [13,14]. In immunologically naïve hosts, immunoglobulin-complexed antigens are efficient stimulators of antibody and cytotoxic T-cell responses [15]. However, most studies using immunoglobulin-complexed antigens have used prototype antigens that have little impact on their intracellular processing. This is not the case for IC-HAdVs. The endosomolytic activity of protein VI, an internal capsid protein, prevents the canonical processing of the IC-HAdVs by enabling the escape of HAdV capsid and its genome from endosomes into the cytoplasm [14]. In the cytoplasm, the HAdV genomes are detected by absent in melanoma 2 (AIM2), a cytosolic pattern recognition receptor (PRR) [16]. AIM2 engagement of the 36 kb HAdV-C5 genome induces pyroptosis, a pro-inflammatory cell death in conventional DCs [17]. Pyroptosis entails inflammasome formation, caspase 1 recruitment/auto-cleavage/activation, pro-IL-1β processing, gasdermin D (GSDMD) cleavage, GSDMD-mediated loss of cell membrane integrity, and IL-1β release [18,19].

Just as immune responses need to be initiated, suppression of cellular responses are primordial to avoid excessive tissue damage and feature prominently in acute and chronic infection [20–22]. Control of antigen-specific T cells can be mediated in part by peripherally induced regulatory T cells (T_regs_) [23]. When regulation of immune responses goes awry, antigen-specific T_regs_ can favor the establishment of persistent viral infections. Moreover, tolerogenic DCs are required for antigen-specific T_reg_ formation. The variable phenotype and functionality of tolerogenic DCs are globally characterized by a semi-mature profile encompassing cell surface costimulatory molecules, cytokine expression and secretion, and antigen uptake and processing [24,25].

The goals of our studies were to determine how HAdV-specific humoral immunity impacts the cellular response to HAdVs, and whether this might affect persistence. Initially, we found that healthy adults harbor HAdV-specific T_regs_. We then demonstrated that IC-HAdV5-challenged human DCs induce a tolerogenic phenotype in bystander DCs. We show that the bystander DCs are capable of taking up and presenting HAdV antigens, and can drive naïve T cells to mature into HAdV-specific T_regs_. Our study reveals a mechanism by which an antiviral humoral responses could, counterintuitively, favor virus persistence.

## Results

### HAdV-specific T_regs_ in healthy donors dampen HAdV-specific T cell proliferation

Initially, we asked if healthy adults harbor HAdV-specific T_regs_ and if so, are they capable of dampening HAdV-specific T-cell proliferation. To address these questions, we prescreened a cohort of healthy individuals using an IFN-γ ELISpot assay for a memory T-cell response to HAdV5 using a pool of overlapping hexon peptides (hexon is the major protein in the HAdV capsid). PBMCs from individuals with a spot forming unit ratio 5-fold greater than mock-treated cells were selected for further analyses. Because inducible T_regs_ can produce IL-10 in response to their cognate antigen, the ability of HAdV-specific CD4^+^ T cells to produce IL-10 as well as IFN-γ, TNF, and IL-2 was assessed by multi-parametric flow cytometry. Consistent with our previous results [13], the cytokine profile of HAdV-specific memory CD4^+^ T cells was dominated by polyfunctional IFN-γ^+^/IL-2^+^/TNF^+^/IL-10^−^ cells (approximately 25% of total HAdV-specific CD4^+^ T cells) and IFN-γ^+^/IL-2^-^/TNF^-^/IL-10^−^ cells (approximately 20%) (**Figure 1A**, a representative donor). We then characterized the combinations of the responses and the percentage of functionally distinct populations in all donors (**Figure 1B**). Each slice of the pie chart corresponds to HAdV-specific CD4^+^ T cells with a given number of functions, within the responding T-cell population. Of note, IL-10-producing HAdV-specific CD4^+^ T cells, which were approximately 5% of total, were predominantly IFN-γ^-^/IL-2^-^/TNF^-^. To determine if the IL-10 producing T cells have a T_reg_ phenotype, the expression of conventional T_reg_ markers, CD45RO, CD25, FoxP3, and CD127 [26], were assessed. We found that approximately 8% of the IL-10 producing T cells were CD25^+^/FoxP3^+^/CD127^dim^. By contrast, most of IFN-γ producing HAdV-specific CD4^+^ T cells harbored a conventional memory phenotype (CD45RO^+^/FoxP3^-^/CD25^-^ /CD127^+^) (**Figure 1C**). These data demonstrate the presence of HAdV-specific T_regs_ in healthy adults.

**Figure 1).**
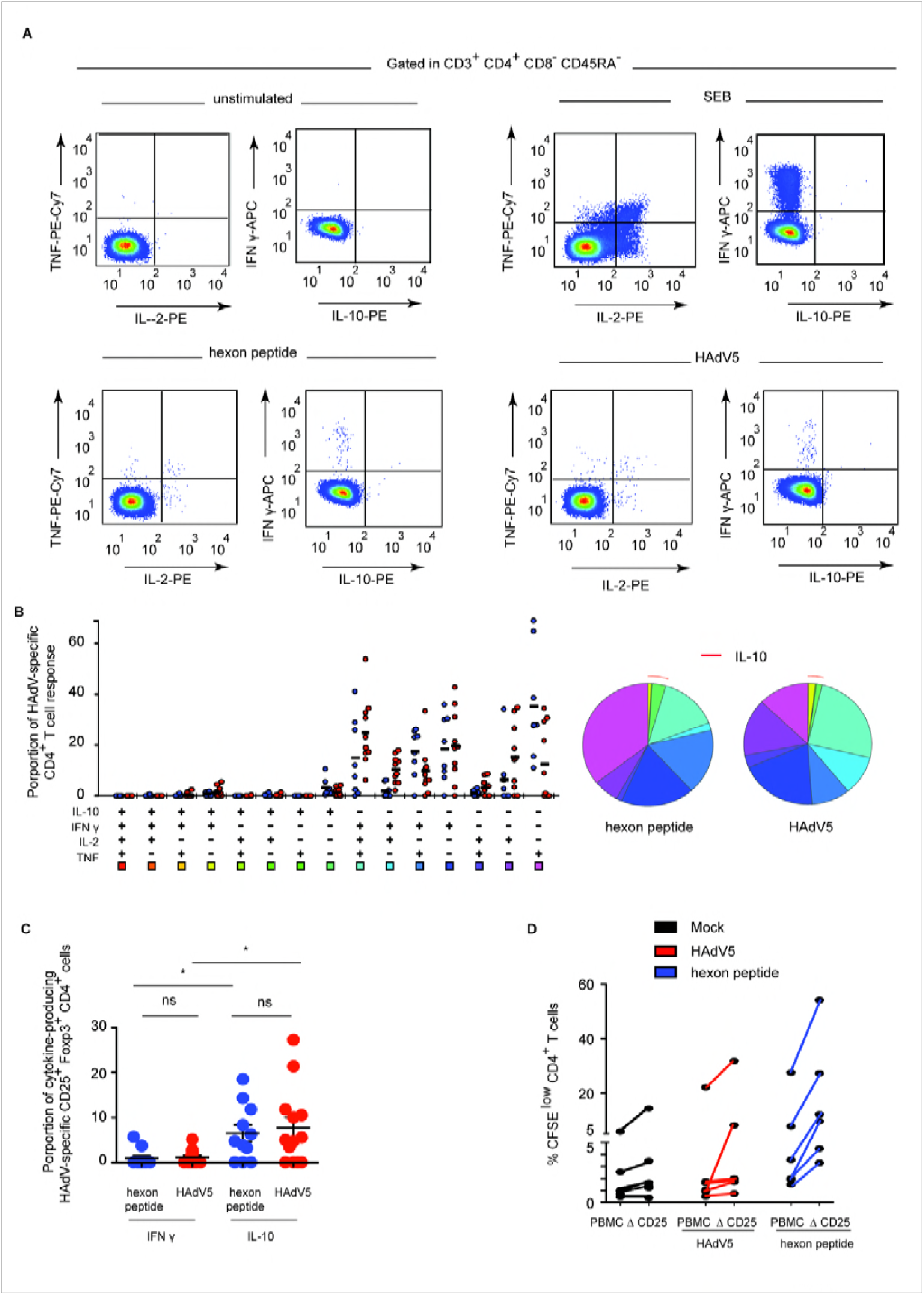
HAdV-specific T_regs_ are present in normal healthy adults. **A**) Representative flow cytometry profile of hexon peptides- and HAdV5-specific (bottom left and right panels) CD4 T cells producing TNF, IL-2, IFN-γ, and IL-10 in a representative subject. Top left panel: mock stimulated (negative control). Top right panel: cytokine profiles of CD4 T cells stimulated with SEB (Staphylococcal enterotoxin B, positive control). **B**) Cumulative (n = 11 donors) cytokine profiles of hexon peptides (blue points) and HAdV5 (red points) CD4 T cells producing TNF, IL-2, IFN-γ, and IL-10. All possible combinations of responses are shown on the x-axis, and the percentage of functionally distinct cell populations within the CD4 T-cell populations are shown on the y-axis. Responses are grouped and color-coded on the basis of the number of functions. The pie chart summarizes the data, and each slice corresponds to the fraction of CD4 T cells with a given number of functions, within the responding CD4 T cells. Red arcs correspond to IL-10 producing CD4 T cell. **C)** Proportion of IL-10 or IFN-γ-producing HAdV5-specific CD4 T cells expressing CD25 and FoxP3 among IL-10, or IFN-γ-producing HAdV5-specific CD4 T cells. **D)** Proliferation of CFSE-labeled PBMCs and PBMCs depleted in CD25^+^ cells (ΔCD25) activated by HAdV5 (red lines) or hexon peptides (blue lines) and cultured for 7 days. The cells were analyzed by flow cytometry for proliferation (CFSE^low^) and CD4 using FlowJo software (n = 6 donors and assayed in duplicate).

To determine if putative HAdV-specific T_regs_ have regulatory functions, we incubated CFSE-labeled PBMCs, or PBMCs depleted in CD25-expressing cells, with a hexon peptide pool and quantified T-cell proliferation. We found that depletion of all CD25^+^ cells caused CD4^+^ cells to proliferate greater than control peptide-challenged CD4^+^ cells **(Figure 1D)**, suggesting that the HAdV-specific T_regs_ in the CD25^+^ population can restrict the proliferative anti-HAdV-specific T cells. Taken together, these data indicate that a fraction of HAdV-specific CD4^+^ T cells harbors an inducible T_reg_ phenotype, and that healthy adults have T_regs_ that dampen the proliferation of HAdV-specific T cells.

### Phenotypic maturation of bystander DCs

A prerequisite for antigen-specific T_reg_ formation is the presence of antigen-presenting tolerogenic DCs [27,28]. Because the cellular profile of HAdV5-challenged DC [29] is inconsistent with that of tolerogenic DCs [29], we asked if IC-HAdV5 could be involved in the generation of HAdV-presenting tolerogenic DCs. When HAdV5 is mixed with neutralizing antibodies from human sera, 200 nm-diameter complexes are formed that induce DCs to undergo pyroptosis, or, if the DC does not die, a hypermature profile [13,14]. As these profiles are also inconsistent with that of tolerogenic DCs, we hypothesized that it was not due to IC-activated DCs, but rather an effect on bystander DCs.

To assess the impact of IC-HAdV5-induced pyroptosis and DC maturation on bystander DCs we developed a transwell assay (**Figure S1A** for schematic). Briefly, CD14^+^ monocytes isolated from fresh buffy coats were induced to differentiate into immature DCs for 6 days. Immature DCs seeded in 12-well plates were mock-treated, challenged with bacterial lipopolysaccharides (LPS), HAdV5, IgGs, or IC-HAdV5 (these cells will be referred to “direct DCs”). At 6 h post-challenge, a transwell insert was added and naive immature DCs (bystander DCs) from the same donor were seeded in the upper chamber (see **Figure S1B-D** for controls concerning transfer of HAdV5 particles between chambers and cell death). Twelve hours after adding the bystander DCs to the upper chamber, the direct and bystander DCs were collected and assayed as described below. Compared to bystander DCs stimulated by direct DCs challenged with IgG or HAdV5, bystander DCs stimulated by IC-HAdV5-challenged DCs increased their cell surface levels of the maturation/activation markers CD80, CD83, CD86 (**Figure 2A)**, CD40, and MHC II (**Figure S2A**). The level of CD86 on bystander DCs tended to increase as the number of IC-HAdV5 particles increased during the stimulation of the direct DCs (**Figure S2B**). The cell surface increase of CD86 and CD83 was also accompanied by an increase in total (cell surface + intracellular) CD86 and CD83 levels (**Figure 2B**). Together, these data demonstrate that IC-HAdV5-challenged DCs enhanced the synthesis and cell surface expression of maturation/activation markers on bystander DCs.

**Figure 2).**
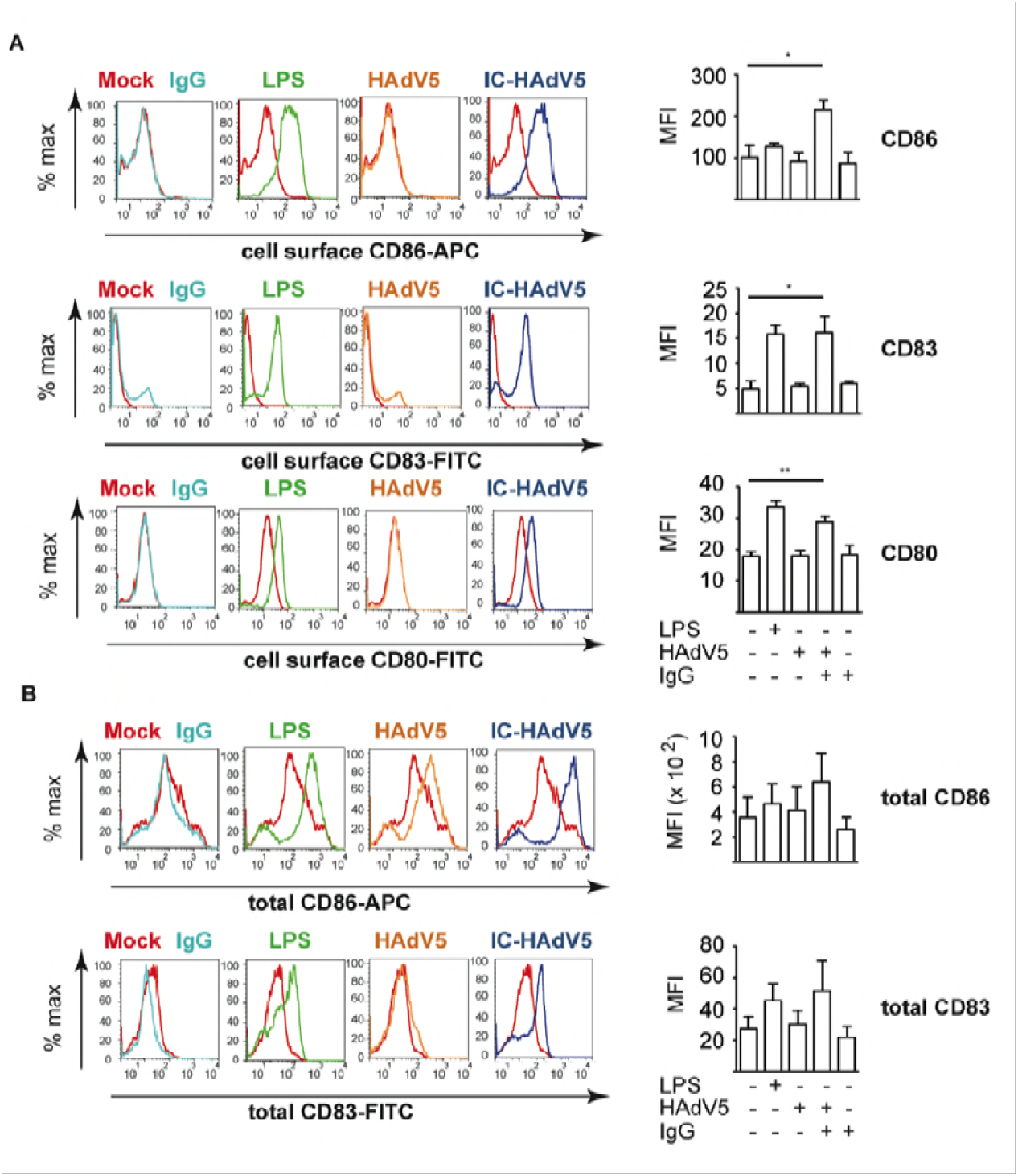
Activation/maturation marker expression in bystander DCs. Representative flow cytometry profile of cells that were mock-treated (red), challenged with IgG (light blue), LPS (green), HAdV5 (orange), or IC-HAdV5 (dark blue). **A)** The cell surface expression of CD86 (top panels), CD83 (middle panels), and CD80 (bottom panels) were quantified 12 h post-stimulation. The profile of mock-treated cells is included in each panel as a reference. Assays were carried out in duplicate in 5 donors with similar results. **B)** Representative flow cytometry profile of total (intracellular and extracellular) CD86 and CD83 levels in bystander DCs incubated with DCs challenged with IgG, LPS, HAdV5, or IC-HAdV5 12 h post-stimulation (color-coded as in **A**). The mock-treated cells are included in each panel as references. The graphs to the right are the cumulative data from 4 donors and performed in duplicate. Error bars are ± SEM. * *p* < 0.05, ** *p* < 0.01.

### The cytokine transcriptome of bystander DCs suggests a tolerogenic profile

To characterize bystander DC functional capabilities we used an 84-plex inflammatory cytokine, chemokine and their receptor mRNA array to quantify transcriptional changes (**Figure S2C** for the list of mRNAs that gave unique amplification profiles). Stimulation of bystander DCs with the milieu from HAdV5-challenged DCs (without IgGs) led to notable increases (>50 fold) in mRNA levels of Th1/Th17 cell activation/differentiation markers (e.g. *CXCL9, CXC10 & CXC11*) (see **Figure S2C** for all data and **Figure 3A** left hand columns for selected data). By contrast, the bystander DC response to the IC-HAdV5-challenged DCs was greater with respect to the number of mRNAs altered (>20) and magnitude (up to 10,000-fold increase) (**Figure 3A** right column, and **Figure S2C** middle column). Of particular relevance was the lack of *TNF* mRNA by bystander DC because tolerogenic DCs should not, *a priori*, secrete TNF. To better understand the transcriptional responses of the different conditions, we applied a principal component analysis (PCA) to find patterns in these data sets. We found that two principal components (see **Materials & Methods** for genes in the F1, F2, and F3 axes) explained 89% and 39% of the total information, respectively, and each stimulus is distinguishable from the others (**Figure 3B**).

**Figure 3).**
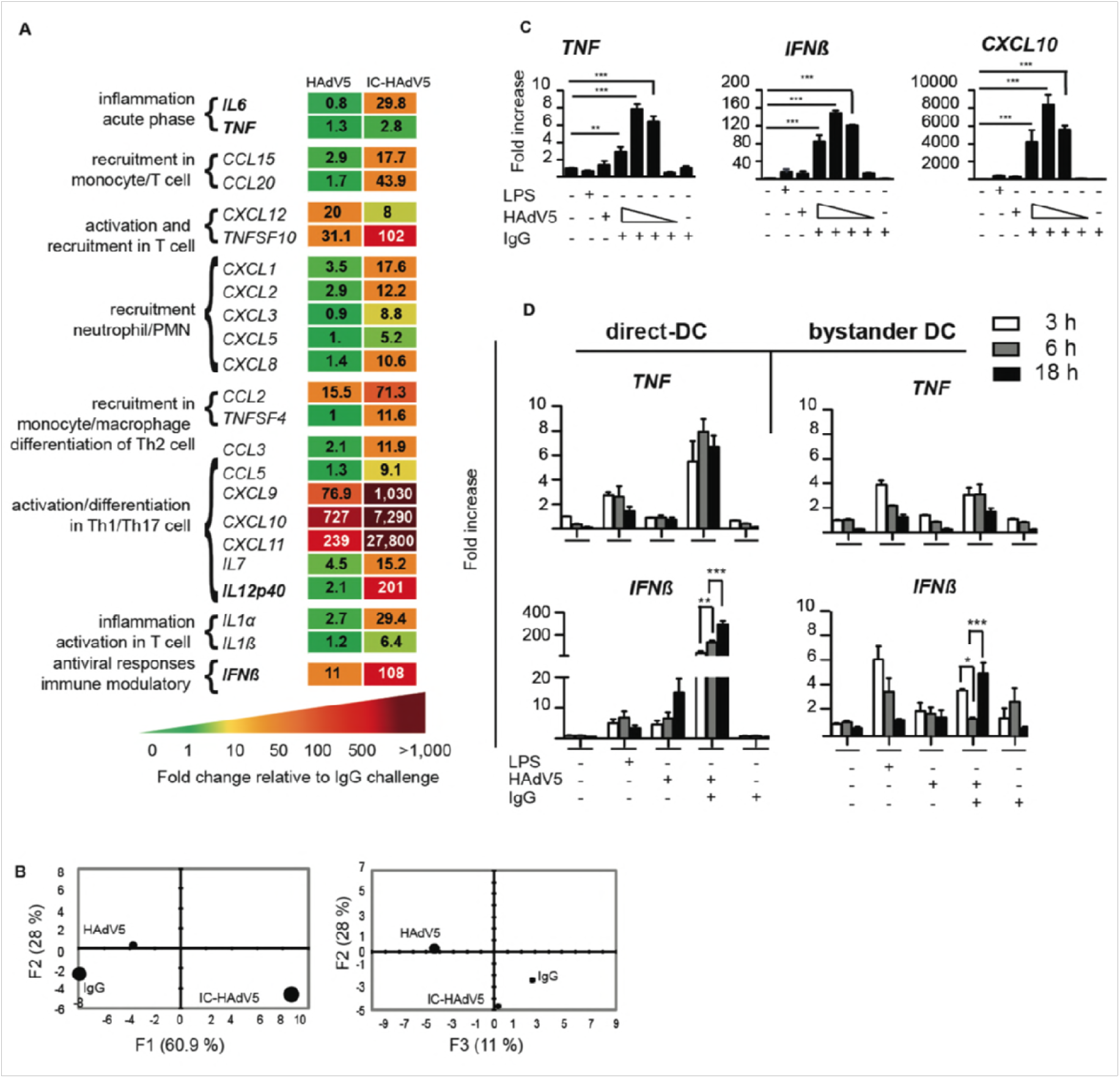
Bystander DC cytokine transcription profile. **A)** Transcription profile of selected cytokines from bystander DCs at 12 h post-activation following exposure to HAdV5- or IC-HAdV5-challenged direct DCs. The transcription profile of bystander DCs created by IgG-challenged direct DCs was used as the baseline. For the genes in bold, primer sequences were designed in-house (n = 2 donors). **B)** Principal component analysis (PCA) of the changes in the 66 mRNAs included in the array. Three principal components showed 61% (F1), 28% (F2), and 11% (F3) accordance. **C)** *TNF, IFNβ*, and *CXCL10* mRNA levels in bystander DCs. Direct DCs were mock-, IgG-, LPS-, HAdV5-, or IC-HAdV-challenged. IC-HAdV5s were used at 20 × 10^3^, 10 × 10^3^, 5 × 10^3^, or 1 × 10^3^ physical particles/direct DC. Assays were carried out in 3 donors in at least duplicates. The fold increase is shown as mean ± SEM. See **Table S1** for additional statistical analyses. **D)** Kinetics (3, 6, 18 h) of *TNF (top panels)* and *IFNβ* (bottom panels) mRNA levels of THP-1-derived DC challenged with LPS, HAd5, IgG or IC-HAdV5 (left panels) or bystander DCs (using the milieu from the direct DCs in the left panel) (right panels). Three independent assays in duplicate were performed. The fold increase is shown as the mean ± SEM. *P* values were derived from one-way ANOVA with Dunnett’s test: ** *p* < 0.01 and *** *p* < 0.001.

Because a cell infected by one HAdV particle could produce >10^4^ virions ∼36 h later, local and global HAdV levels, as well as IC-HAdV formation, are dynamic. Of note, IC-HAdV5 causes a dose-dependent induction of pyroptosis in direct DCs [14]. We therefore extended the mRNA array analyses by quantifying dose-dependent response of bystander DCs. Using RT-qPCR we analyzed *TNF, IFNβ* and *CXCL10* (**Figure 3C**) and *IL1β, IL12* (p40), *CCL3* and *IL6* (**Figure S2D)** mRNA levels. In all cases the transcriptional response of bystander DCs varied depending on the IC-HAdV5 challenge dose. These data suggest that the bystander DC response is linked to the percentage of direct DCs undergoing pyroptosis [14].

To characterize time-dependent transcriptional changes in direct DC and bystander DCs, we compared mRNA levels of *TNF, IFNβ* (**Figure 3D**), *Mip-1α* and *IL6* (**Figure S3,** which also includes dose-dependent response). Globally, mRNAs that code for pro-inflammatory molecules were 2 to 10-fold greater in direct DCs than in bystander DCs. In addition, only *IL1β* and *Mip-1α* mRNA levels changed significantly (*p <* 0.01) over time. These data demonstrate that bystander DCs have a semi-mature tolerogenic transcriptional profile, which is linked to DC pyroptosis, and lack noteworthy levels of *TNF* mRNA [30].

### Cytokine secretion by bystander DCs is consistent with a tolerogenic profile

To examine the events downstream the transcriptional response, we quantified the secreted cytokine from direct and bystander DCs. Because proteins can readily diffuse across the transwell membranes, bystander DCs were removed from the upper chamber 12 h post-challenge, rinsed, and then placed in a separate well with fresh medium for 9 h before collecting the medium. The direct DC medium was collected at 12 h post-challenge, or after a wash at 12 h and then collected 9 h later (21 h) to compare conditions similar to that used for bystander DCs (see **Figure 4A** for schematic). Challenging DCs with HAdV5 alone had a modest effect on their secretome with the exception of a 5 to 10-fold increase in TNFSF10, and CXCL9 & 10 levels (**Figure 4B,** second column from the left). By contrast, IC-HAdV5-challenged DCs responded with increases of >15 fold in approximately half of the cytokines (**Figure 4B,** middle columns). These data are consistent with previous results showing the robust maturation of IC-HAdV5-challenged DCs [13,14]. HAdV5-challenge DCs that were rinsed 12 h post-stimulation had overall lower cytokine levels than prior to washing, but TNFSF10, CXCL9, CXCL11, and CCL5 levels remained robust (**Figure 4C,** middle columns). Interestingly, instead of a positive correlation between the cytokine secretion and the IC-HAdV5 dose, we found that as the IC-HAdV5 dose increased, the cytokines secreted by direct DCs tended to decrease (**Figure 4C,** middle columns).

**Figure 4).**
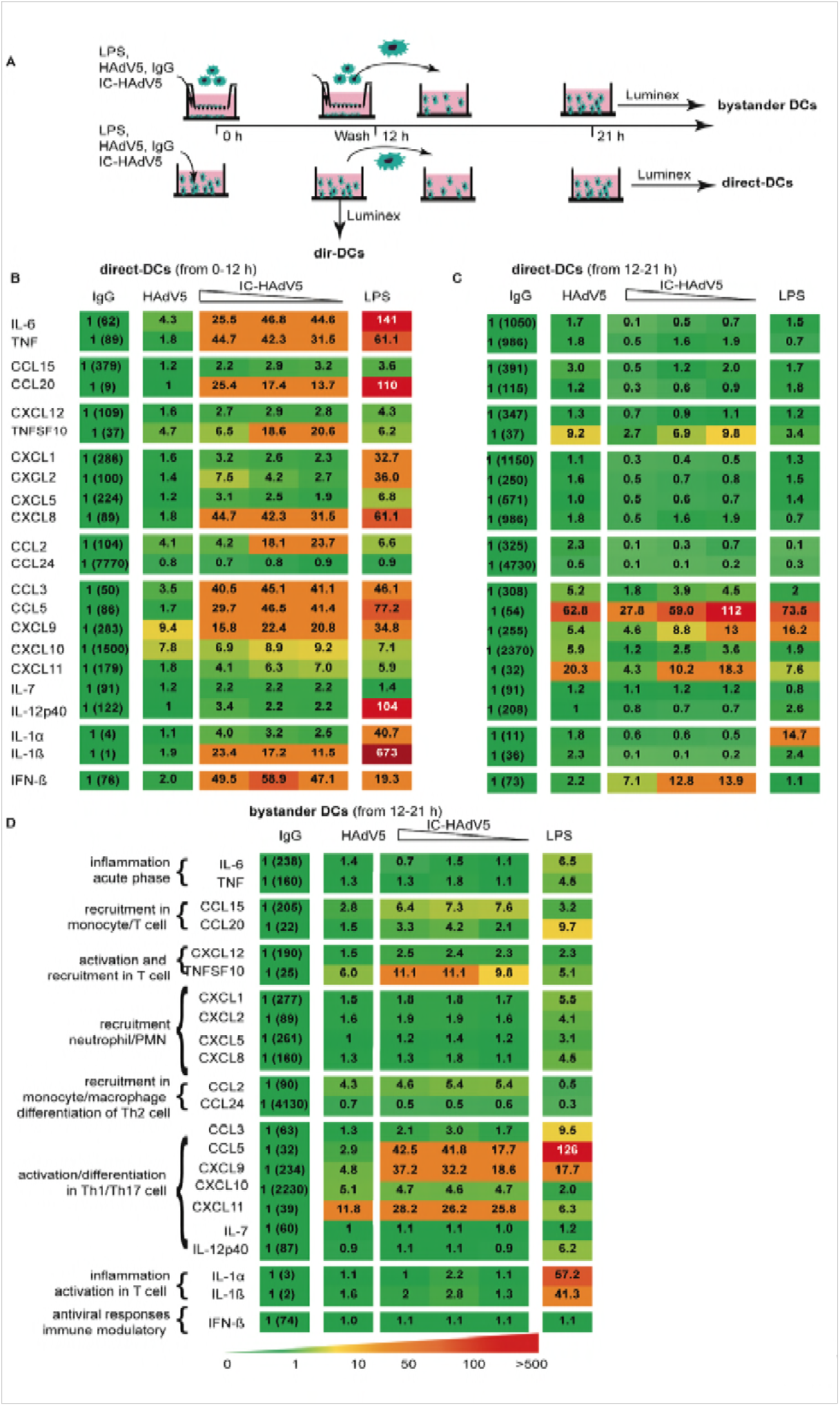
Direct and bystander DC cytokine secretomes. **A)** Schema showing how DCs were activated with LPS, HAdV5, IgG, and IC-HAdV5 and when the cell supernatants were harvested. **B)** Cytokine secretion from IgG-, HAdV5-, dose-dependent IC-HAdV5-, and LPS-challenged DCs at 12 h. **C)** Cytokine secretion from IgG-, HAdV5-, dose-dependent IC-HAdV5-, and LPS-challenged DCs at 12-21 h. **D)** Cytokine secretion from bystander DCs (the stimulus used to challenge the direct DCs is above each column at 12-21 h). The color code shows relative increases compared to IgG-challenged-DCs (raw values in first column). The assays were performed twice in duplicate with similar results.

Using HAdV5-challenged DCs (without IgGs) to generate bystander DCs, we found that the latter secreted 3 to 12-fold higher levels of TNFSF10, CCL5, CXCL9, CXCL10 and CXCL11 compared to bystander DCs exposed to the medium from IgG-challenged DCs (**Figure 4D,** second column). Similarly, when bystander DCs were generated using IC-HAdV5-challenged DCs, the level of the above five cytokines also increased. In addition, three chemokines involved in immune cell recruitment (CCL15, CCL20, and CCL2) increased >3 fold. Consistent with the transcriptome analyses, we did not find a notable dose-dependent effect on bystander DCs when direct DCs were incubated with increasing IC-HAdV5 particles (**Figure 4D,** middle columns).

Together, these data suggest that the release of pathogen-associated molecular patterns (PAMPs), danger-associated molecular patterns (DAMPs), and/or the increased levels of cytokines secreted by a greater number of DCs that do not undergo pyroptosis, are key factors in bystander DC maturation. In addition, the environment created by IC-HAdV5 induces a semi-mature cytokine secretion profile in bystander DCs.

### Cytokines and pyroptosis-associated factors impact bystander DC phenotype

To determine how cytokines and pyroptosis impact bystander DCs, we used a combination of drugs and mutant viruses to selectively modify the environment created by IC-HAdV5-challenged DCs. To determine the impact of IL-1β, direct DCs were pre-treated with ZVAD, a pan-caspase inhibitor that blocks caspase 1 auto-cleavage and pro-IL-1β processing. Importantly, ZVAD has no effect on TNF and canonical protein secretion **(Figure S4A** and reference [14]). We found that blocking IL-1β production by direct DCs reduced bystander DC maturation as demonstrated by their lower levels of CD86 and CD83 (**Figure 5A-B**). We then used brefeldin A to block ER to Golgi-mediated cytokine secretion in direct DCs (see **Figure S4B** for controls). Of note IL-1β release is not significantly affected by brefeldin A, (**Figure S4C**). In brefeldin A-treated IC-HAdV5-challenged DCs the levels of CD83 and CD86 did not change markedly (**Figure 5C**), while the bystander DCs responded with lower levels of CD83 and CD86 (**Figure 5D)**.

**Figure 5).**
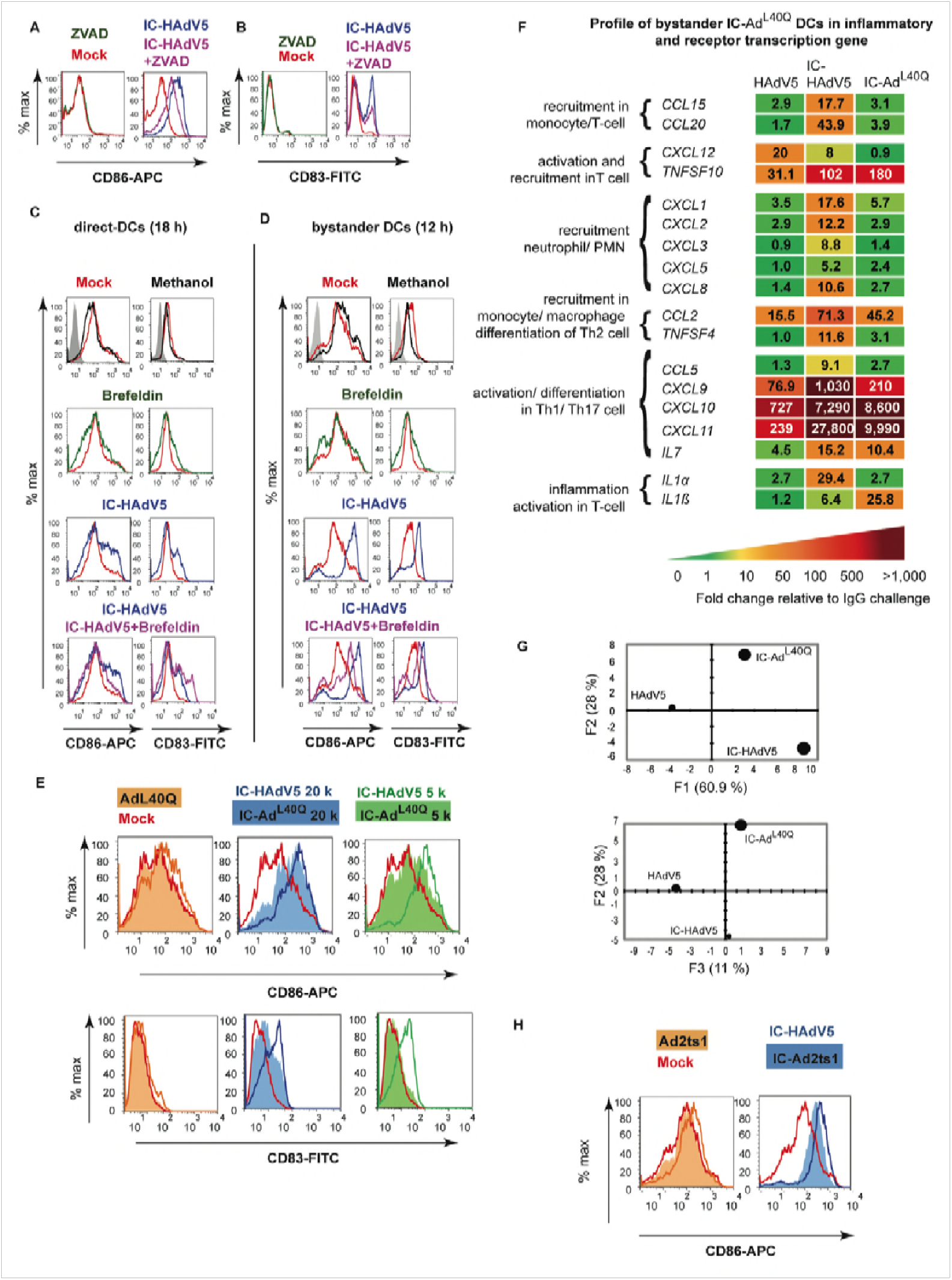
Impact of cytokines and pyroptosis-associated factors on bystander DC maturation. **A & B)** Shown are representative flow cytometry profile of direct DCs pretreated with ZVAD using the approach described in **Figure S1**. The cell surface levels of CD86 and CD83 on bystander DCs was quantified by flow cytometry (n ≥3 donors). **C & D)** Representative flow cytometry profile of IC-HAdV5-challenged DCs treated with brefeldin A and the cell surface levels of CD86 and CD83 quantified by flow cytometry in direct DCs and bystander DCs (n ≥ 5 donors). **E)** Bystander DCs were incubated with milieu generated by DC challenged with increasing doses of IC-Ad^L40Q^ (representative flow cytometry profiles of CD86 and CD83 levels). **F)** Bystander DC inflammatory cytokine mRNA levels were measured by RT-qPCR array following incubation with DC challenged with HAdV5, IC-HAdV5 and IC-Ad^L40Q^. The heat map denotes the fold change relative to DCs challenged by IgGs (n = 3 donors). **G)** PCA of the changes in the 66 mRNAs included in the array when including IC-Ad^L40Q^-. **H)** Bystander DCs were incubated with milieu generated by DCs challenged with increasing doses of IC-Ad2ts1. Representative flow cytometry profiles of cell surface level of CD86 (n = 2 donors).

Next, we generated ICs using Ad^L40Q^ [31], an HAdV5 capsid containing a mutated protein VI that attenuates endosomolysis. While IC-Ad^L40Q^ poorly induces pyroptosis in direct DCs [14], they secrete levels of TNF that are similar to IC-HAdV5-challenged DCs. Furthermore, *IFNβ* and *IL1β* mRNA levels are lower [14]. We found notably lower levels of CD86 and CD83 on bystander DCs following stimulation with the response from IC-HAdV5 versus IC-Ad^L40Q^-challenged DCs. In addition, the reduced maturation/activation effects were only modestly altered by increasing the IC-Ad^L40Q^ dose (**Figure 5E**). Together, these data demonstrate a role for pyroptosis-associated factors in the maturation of bystander DCs.

We then compared cytokine mRNA levels in bystander DCs stimulated by HAdV5-, IC-Ad^L40Q^--, or IC-HAdV5-challenged DCs (**Figure 5F** and **Figure S5A-C**). Consistent with the phenotype, the transcriptional responses of bystander DCs to both ICs were globally higher than to HAdV5 alone. The bystander DC transcriptional response to IC-Ad^L40Q^-challenged DC milieu was generally lower than in IC-HAdV5-challenged DCs, and it was qualitatively distinguishable as determined by PCA (**Figure 5G**).

We then assessed the effect of pyroptosis using IC-Ad2ts1. Ad2ts1 has a hyper-stable capsid due to a mutation in protease that results in failure to process the capsid preprotein [32,33]. We previously showed that IC-Ad2ts1 poorly induces DC pyroptosis, likely because the HAdV genome does not escape from the capsid and therefore does not nucleate AIM2 (see reference [14] and **Figure S5D-F** for Ad2ts1 controls). Of note, TNF levels are comparable in DCs challenged with IC-Ad2ts1 or IC-HAdV5 [14]. Here, we found that IC-Ad2ts1-challenged DC induced an increase of CD86 cell surface levels on bystander DCs (**Figure 5H).** Together, these data demonstrate that cytokines and pyroptosis-associated factors play a role in the activation and semi-maturation of bystander DCs.

### Engagement of TLR4 on bystander DCs

To characterize how bystander DCs are activated, we focused on Toll-like receptor 4 (TLR4). TLR4 is a multifunctional cell surface PRR that can directly or indirectly (by forming a complex with MD-2, CD14, or other PRRs) be activated by extracellular viral components (PAMPs) and, under inflammatory conditions, extracellular high-mobility group box 1 and heat shock proteins (DAMPs) [34–36]. Of note, MD-2 acts as a coreceptor for recognition of both exogenous and endogenous ligands [37–40]. While TLR4 does not bind to, or become activated by, HAdV5 alone [41], TLR4 might be activated by PAMPs or DAMPs that interact directly with the HAdV5 capsid. We therefore used TAK-242 to disrupt TLR4 signaling in bystander DCs (see **Figure S6** for TAK-242 control). As readouts, we used the upregulation of *TNF* and *IL1β* mRNAs, and activation/maturation cell surface markers. When TLR4 signaling was blocked in bystander DCs stimulated by the IC-HAdV5-challenged DC milieu, there was a significant (*p* < 0.05) decrease in *IL1β* mRNA levels and 2-fold decrease of *TNF* mRNA (**Figure 6A**). CD83 and, to a lesser extent, CD86 levels were also reduced (**Figure 6B**). These data suggest that bystander DCs use TLR4 to detect PAMPs and DAMPs released by IC-HAdV5-challenged DCs, leading to changes in bystander DC maturation.

**Figure 6).**
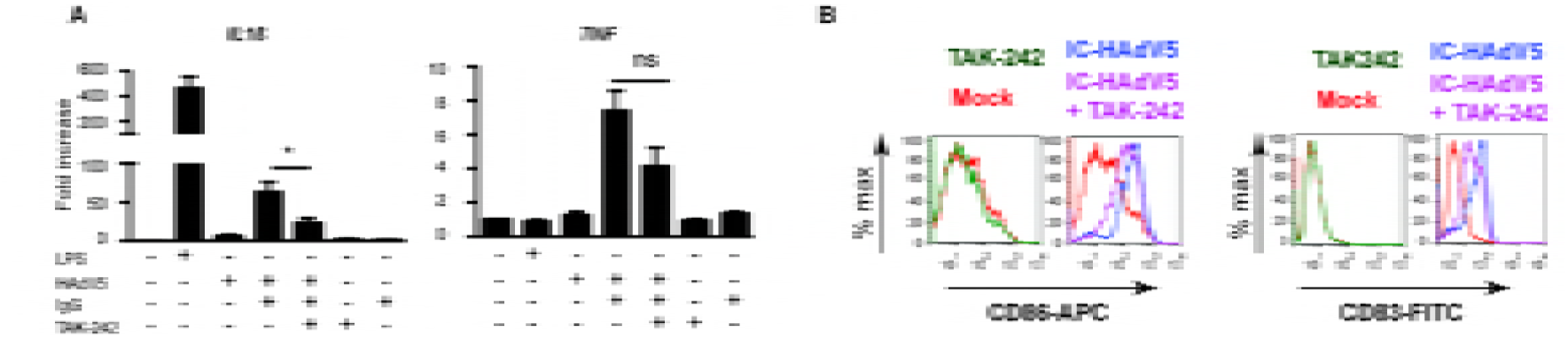
Impact of TLR4 engagement on cytokine transcription and activation/maturation markers in bystander DCs. Involvement of TLR4 signaling in bystander DCs was assessed by **A)** *IL1β* and *TNF* mRNA levels in bystander DCs pre-treated with TAK-242, and then added to milieu of DCs challenged with LPS, IgG, HAdV5 and IC-HAdV5. Fold increase is represented as mean ± SEM (n = 3 donors, * denotes *p <* 0.05) **B)** Representative flow cytometry profiles of CD86 and CD83 cell surface levels in bystander DCs. DCs were mock-treated (red line), TAK-242 alone (with TAK-242 and without direct DCs, green line) challenged with IC-HAdV5 (dark blue) or pretreated with TAK-242 and challenged with the milieu generated from IC-HAdV5-challenged DCs (violet line).

### Minimal loss of phagocytosis in bystander DCs is consistent with tolerogenic profile

Immature DCs survey the extracellular environment by random phagocytosis. Once PRRs are engaged, DC maturation is accompanied by decreased uptake of fluid phase molecules [42]. Of note, a functional hallmark of tolerogenic DCs is their ability to retain some antigen uptake properties. To address the functional maturation of IC-HAdV5-challenged DCs and bystander DCs, we incubated cells with FITC-labeled dextran and quantified uptake by flow cytometry. We found that phagocytosis was modestly downregulated in direct DCs stimulated with HAdV5 or LPS (**Figure 7A**). By contrast, IC-HAdV5-challenged DC phagocytosis was near background levels, consistent with complete maturation (see **Figure S7** for controls) [29]. While bystander DCs had reduced phagocytosis when created by IC-HAdV5-challenged DCs, the bystander DCs still took up 17-fold more FITC-dextran than background levels (**Figure 7B**). These functional data are consistent with semi-mature, tolerogenic DCs.

**Figure 7).**
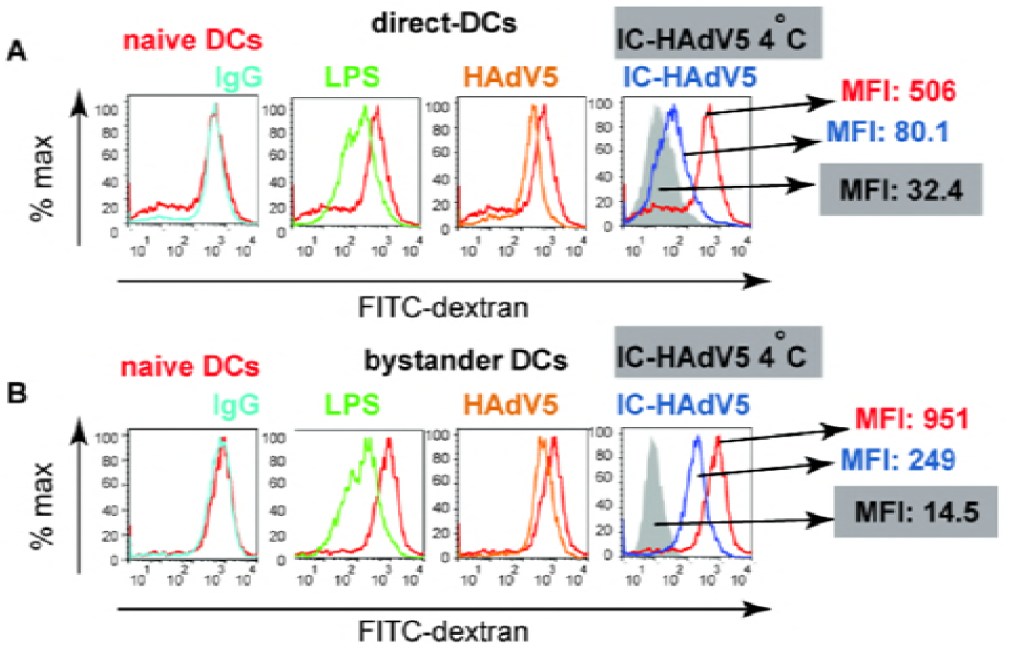
Fluid phase uptake by direct and bystander DCs. Fluid phase antigen uptake by direct and bystander DCs was quantified using FITC-labeled dextran and flow cytometry. **A)** Representative flow cytometry profiles of direct DCs challenged with LPS, HAdV5, IgG, or IC-HAdV5 and **B)** Representative flow cytometry profiles of bystander DCs with the corresponding direct DC milieu. Experiments were performed 3 times, in duplicate, with similar results. Nonspecific binding of dextran to cells was controlled by incubation at 4°C (**Figure S7**). MFI – median fluorescent index.

### Bystander DCs create loop to recruit antigen-presenting cells

While tolerogenic DCs can induce, recruit, and maintain T_reg_ homeostasis, tolerogenic DCs can also create a feedback loop to promote their own generation [43]. Because monocytes are recruited to sites of inflammation [44,45], we compared the functional recruitment capabilities of direct DCs and bystander DCs (see **Figure S8** for setup and controls). Unexpectedly, we found that IC-HAdV5-challenged DCs inhibited monocyte recruitment in an IC-HAdV5 dose-dependent manner (**Figure 8A & B**). Of note, the inhibition was abrogated when the IC-HAdV5-challenged DC were washed, suggesting that inhibitory factors were generated <3 h post-IC-HAdV5 challenge (**Figure 8B**). To determine if pyroptosis-related factors are responsible for the inhibition of monocyte recruitment, we used IC-Ad^L40Q-^ and ZVAD to reduce pyroptosis and IL-1β secretion. The effect of ZVAD was modest and did not markedly influence the monocyte recruitment induced by IC-HAdV5-challenged DC, suggesting that pyroptosis related factors (e.g. IL-1β) did not have a role in this process (**Figure 8C & D**). In contrast to the IC-HAdV5-challenged DC response, the IC-Ad^L40Q^-challenged DC response significantly (*p <* 0.05) increased monocyte recruitment, in an IC-Ad^L40Q-^ dose-dependent manner (**Figure 8E & F**). These data demonstrate that pyroptosis factors other than IL-1β inhibit monocyte recruitment. We then examined the ability of bystander DCs to recruit monocytes. In contrast to IC-HAdV5-challenged DCs, bystander DCs promoted monocyte recruitment (**Figure 8G**). These data are consistent with the bystander DC milieu containing more chemoattractants **(Figure 5**). There was also a trend towards greater recruitment when higher IC-HAdV5 doses were used on the direct DCs.

**Figure 8).**
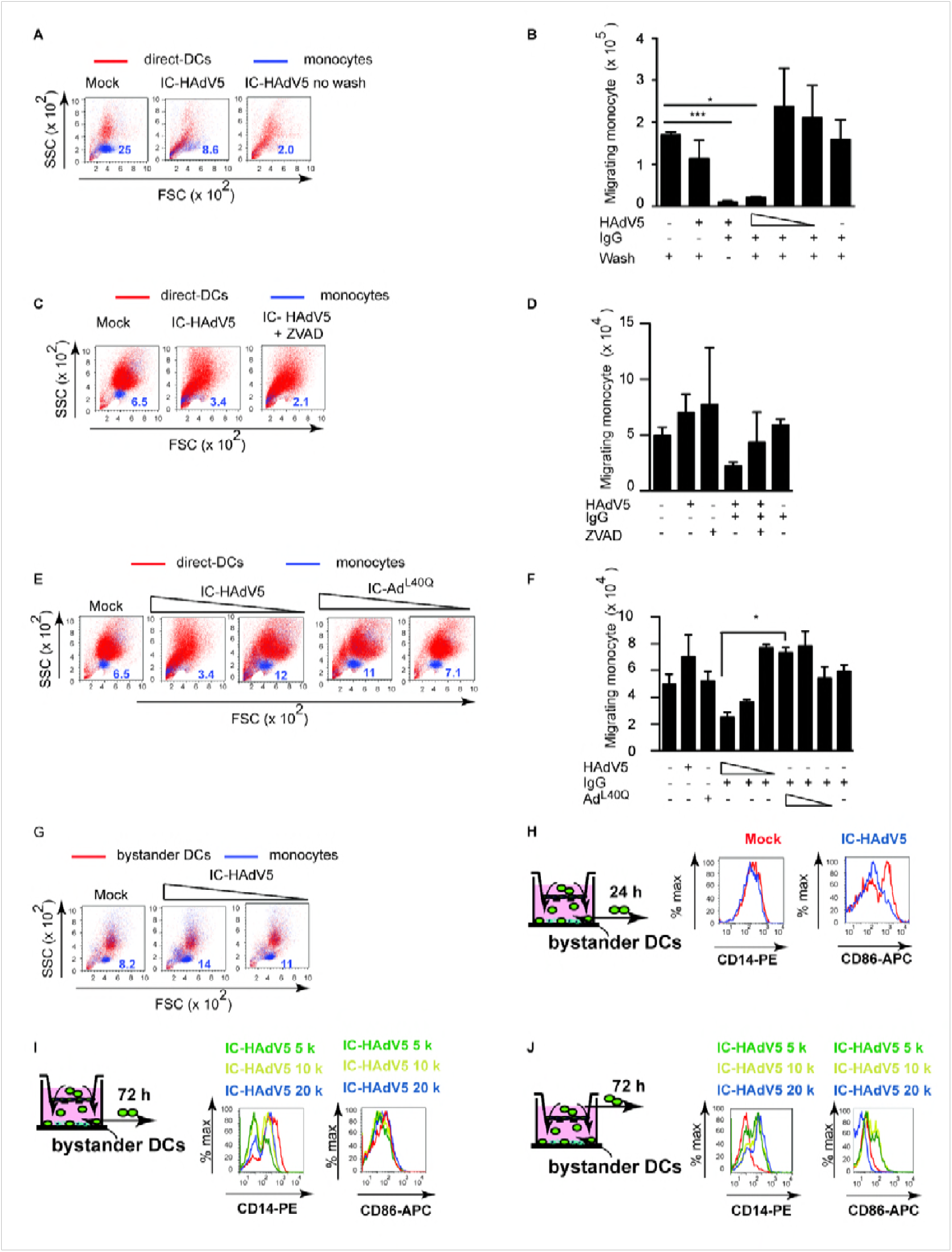
Direct and bystander DC monocyte recruitment and their phenotype. A 5 μm-pore transwell system (see **Figure S8** for details) was used for monocyte migration assays. **A)** Representative FCS/SSC profiles of CFSE-labeled monocyte (blue) that migrated into the lower direct DC chamber; The numbers in dark blue correspond to the percentage of CFSE-labeled monocytes. **B)** Cumulative data from monocyte migration at 24 h into the lower chamber containing DCs challenged with HAdV5, IgGs or decreasing doses (20 × 10^3^, 10 × 10^3^ and 5 ×10^3^) of IC-HAdV5 and washed after 30 min post-challenge; data are mean ± SEM, n = 4 donors; **C)** Representative FCS/SSC profiles of direct DCs (red) pretreated with ZVAD before activation with IC-HAdV5. The CFSE-stained monocytes (blue) were added to the upper compartment and migration was quantified at 24 h; Numbers in dark blue correspond to the percentage of CFSE-labeled monocytes. **D)** Cumulative data from assay in (**C**) n ≥3 donors; **E)** Representative FCS/SSC profiles of DCs (red) challenged with decreasing doses of IC-HAdV5 or IC-Ad^L40Q^ and CFSE-labeled monocyte (blue) recruitment at 24 h. The numbers in dark blue correspond to the percentage of CFSE-labeled monocytes. **F)** Cumulative data from assay in (**E**) using DCs from ≥3 donors; **G)** Bystander DCs activated for 12 h with milieu from DC challenged with increasing concentration of IC-HAdV5. The bystander DCs were seeded in the lower chamber of a transwell and CFSE-labeled monocyte recruitment was quantified by flow cytometry at 24 h (n ≥4 donors). The numbers in dark blue correspond to the percentage of CFSE-labeled monocytes. **H)** Phenotypic characterization of the monocytes (green cells seeded in the upper chamber) recruited by bystander DCs. DCs were challenged with increasing doses of IC-HAdV5 (colored coded on top of each panel). The milieu from these direct DCs was used to generate bystander DCs. The bystander DCs were seeded in the lower chamber of the transwell system and the monocytes that were recruited were characterized for their expression of CD14 and CD86 at 24 h, and **I)** at 72 h. **J)** Monocytes that did not migrate into the lower chamber were also characterized for the CD14 and CD86 levels. The data are representative flow cytometry profiles of experiments carried out in ≥5 donors. *p* values in **B, D,** and **F** were derived from *t*-tests: * *p* < 0.05, ** *p* < 0.01, and *** *p* < 0.001.

Once monocytes migrate into an inflammatory environment they acquire distinct phenotypic and functional profiles [46]. One phenotypic hallmark of monocyte differentiation is CD14, which is high on monocytes and macrophages, but lower on DCs. We therefore characterized migrating and static monocytes for CD14 and CD86 levels at 24 and 72 h (see schematic at the left of each panel in **Figure 8H-J** for the times and location of cells, and **Figure S8** for controls). At 24 h the level of CD14 on monocytes that had migrated into the bystander DCs environment did not change markedly, while CD86 levels were lower (**Figure 8H**). At 72 h the recruited monocytes had two distinct populations based on CD14 levels (**Figure 8I**). The decrease in CD14 levels suggested that they differentiated into DCs, while the CD86 levels suggest the maintenance of an immature phenotype. In addition, monocytes recruited by bystander DCs had increased CD14 levels. By contrast, CD86 levels decreased on monocytes in the upper chamber (bottom chamber containing bystander DCs) (**Figure 8J**).

Together, these data demonstrate that DCs challenged with IC-HAdV5 inhibit monocyte recruitment. Monocytes recruited to the bystander DC environment was abetted by pyroptosis of direct DCs. Recruited monocytes had reduced CD14 levels, possibly due the engagement and internalization of TLR4/CD14 complexes. Monocyte-DC contact also appeared to favor the increase in cell surface levels of activation/maturation markers. We concluded that the dynamic environment created by bystander DCs is consistent with a feed-forward loop to foster tolerogenic DCs.

### Bystander DCs induce memory T-cell proliferation and naïve CD4 T cells towards HAdV-specific T_regs_

A functional characteristic of tolerogenic DCs is that they take up and present antigens. Therefore, we asked if some of the bystander DCs generated in our ex vivo model are capable of inducing proliferation of HAdV5-specific memory T cells. We used IC-HAdV-challenged DC to generate bystander DCs, which were then added to CFSE-labeled PBMCs. Seven days post-incubation we found that CD3^+^/CFSE^low^ cells harbored memory T cell markers (CD45RO^+^/CD45RA^-^) (**Figure 9A**). These data are consistent with the potential of some of the bystander DCs to maintain fluid phase uptake and subsequent presentation of HAdV5 antigens to memory T cells.

**Figure 9).**
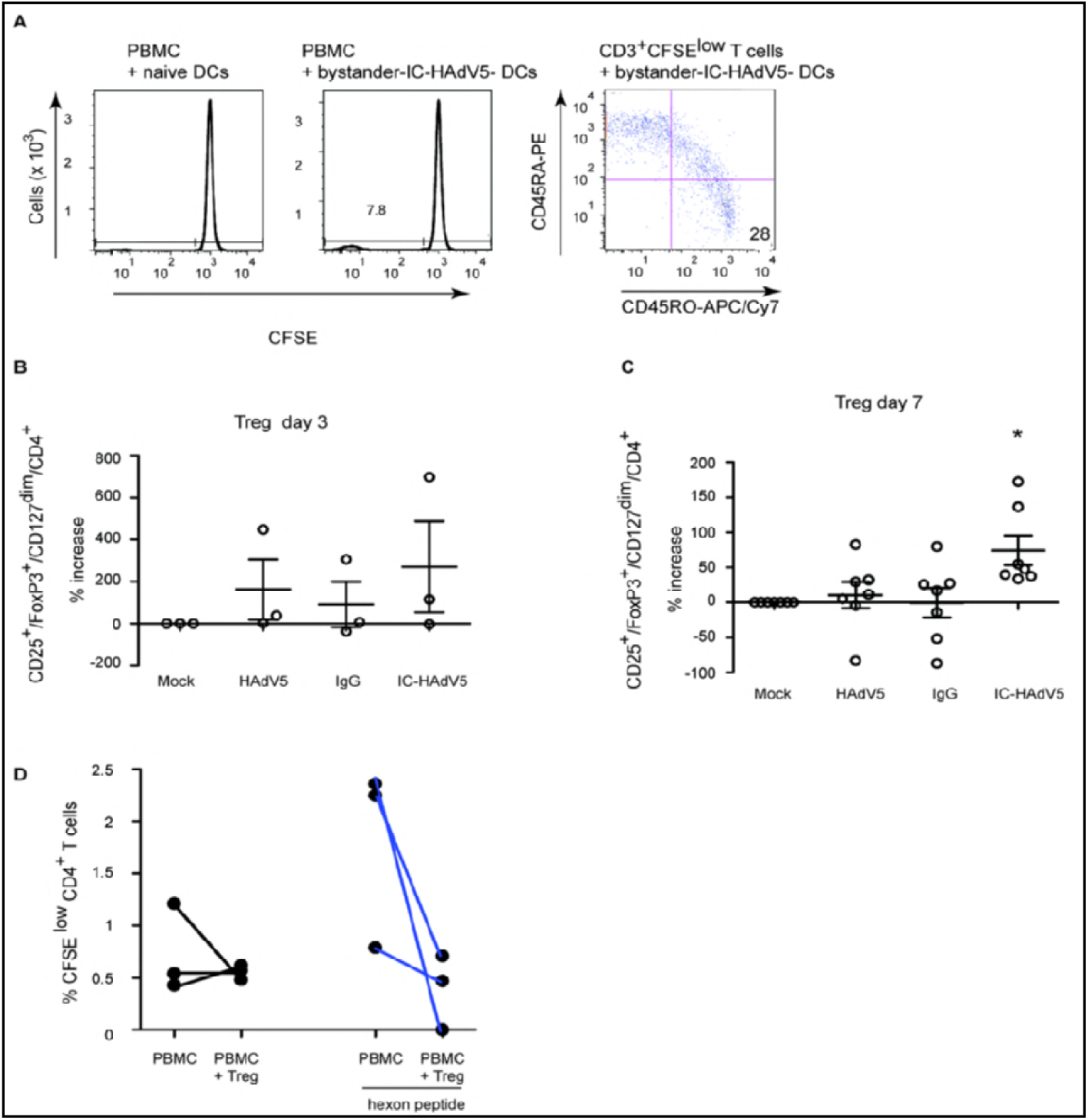
Bystander DCs induce memory T cell proliferation and promote naïve CD4 cells towards T_reg_ phenotype that inhibit proliferation of anti-HAdV T cells. **A)** Bystander DCs, generated via IC-HAdV stimulation of direct DCs or mock-treated DCs, were incubated with CFSE-labeled PBMCs and proliferation was quantified by flow cytometry. CD3^+^/CFSE^low^ cells were screened for memory T cell profile (CD45RO vs. CD45RA). Bystander DCs, generated using the media from DCs challenged with IgG, HAdV5, and IC-HAdV5, were incubated with 10^5^ naive CD4^+^ T (ratio of 10:1 PBMC/ bystander DC) cells isolated from the same donors. The percentage of T_regs_ (CD25^+^/FoxP3^high^/CD127^dim^/CD4^+^ cells) varies between 1 to 5% of CD4^+^ cells in peripheral blood. The number of T_regs_ in the CD4^+^ cell population, ± bystander DCs, was quantified by analyzing 50,000 cells by flow cytometry. The results are presented as percentage of increase of mock-treated cells at day 3 (**B**) or day 7 (**C**). **D)** T_regs_ generated by bystander DCs reduce the proliferation of memory anti-HAdV CD4 T cells. Experiments were carried out in ≥7 donors with similar results. * *p <* 0.05 vs. mock, HAdV5 and IgG.

In addition to antigen presentation, tolerogenic DCs can induce naïve CD4^+^ cells to become T_regs_. To address this functional characteristic, bystander DCs were generated and incubated with autologous naïve CD4^+^/CD45RA^high^ cells for 3 or 7 days. The T cells were then assayed by multi-parametric flow cytometry for CD4, CD25, CD127 and FoxP3, markers that are indicative of T_regs_. While activated T cells transiently express FoxP3 (**Figure S9**), the relatively low-level does not result in acquisition of suppressor activity [27]. By contrast, stable and high levels of FoxP3 can be used to identify *bona fide* T_regs_. At day 3, naïve T cells expressed T_reg_ markers in all conditions (except mock-treated direct DCs) (**Figure 9B**). At day 7, the number cells with T_reg_ phenotype was near background following incubation in the milieu of mock-, IgG-, or HAdV5-challenged direct DC (**Figure 9C**). By contrast, bystander DC created from IC-HAdV-direct DCs had a significant (*p* < 0.05) increase in cells with a T_reg_ profile. This functional assay demonstrates that bystander DCs can induce naïve CD4 into cells with a T_reg_ phenotype, further supporting our conclusion that they are tolerogenic DCs.

As shown in **Figure 1D**, healthy adults harbor CD25^+^ cells can inhibit HAdV-specific CD4^+^ cell proliferation. We therefore asked if the tolerogenic bystander DCs generated in our ex vivo assay could induce the production of HAdV-specific T_regs_. To address this question we isolated PBMCs, CD14^+^ monocytes, and naïve CD4^+^ T cells from 3 donors that harbored anti-HAdV memory T cells (see **Figure S10** for flow chart). Briefly, monocytes were used to create direct DCs that were incubated with IC-HAdV-C5. Bystander DCs were generated as previously described. VPD 450-labeled naïve CD4^+^ T cells were incubated with bystander DCs to generate T_regs_. VPD 450^low^/CD4^+^/CD25^+^ cells (600 to 5,000 cells) were isolated by FACS and mixed with CFSE-labeled PBMCs ± hexon peptides. We found that the ex vivo generated T_regs_ from all donors reduced the proliferation of anti-HAdV T cells (CFSE^low^/CD4^+^) (**Figure 9D**). These data demonstrate that HAdV-specific T_regs_ can be generated via bystander DCs.

## Discussion

HAdV infections lead to multifaceted, robust, long-lived cellular and humoral responses in most young immunocompetent adults. Nonetheless, several HAdV types somehow circumvent immune surveillance to establish persistent infections. It is well documented that HAdV neutralizing antibodies are type specific, while the anti-HAdV cellular response is cross-reactive [1,3,8,47–49]. In addition, it is the anti-HAdV cellular response that protects us from reactivation of persistent infections. The dichotomy between the two arms of the adaptive immune response led us to address how anti-HAdV antibodies influence HAdV-specific T-cell responses. In this study, we initially asked if healthy adults harbor HAdV-specific T_regs_, which would be indicative of a path towards HAdV persistence. We then explored how HAdV-specific tolerogenic DCs and T_regs_ could be generated. We previously showed that IC-HAdV5s are internalized by, and aggregate in, DCs [14]. Following protein VI-dependent endosomal escape of the capsid, the viral genome is engaged by AIM2 in the cytoplasm. AIM2 nucleation induces ASC (apoptosis-associated speck protein containing a caspase activation/recruitment domain) aggregation, inflammasome formation, caspase 1 auto-activation, pro-IL-1β and GSDMD cleavage, and GSDMD-mediated loss of cell membrane integrity. Here we demonstrate that the pyroptotic environment plays a significant role the creation of tolerogenic bystander DCs. We further show that these bystander DCs can induce HAdV5-specific memory T cells to proliferate, and can drive naïve CD4 cells towards a T_reg_ phenotype. The T_regs_ generated in this *ex vivo* assay are capable of inhibiting the proliferation of anti-HAdV T cells. We therefore propose that HAdV neutralizing IgGs [14] abet HAdV persistence.

Our assays using a human pathogen, naturally occurring human antibodies and primary blood-derived human cells address the immune cell-based mechanisms of adenovirus persistence. Yet, *ex vivo* results cannot unequivocally show causality. Host-pathogen-based studies have often used mice to address questions underlying disease-immune relationships. However, the impact of HAdVs on human and mouse DCs is notably different. Furthermore, we are unaware of studies directing addressing the impact of murine adenovirus (MAV) on murine DCs. In 1964, D. Ginder showed that a MAV can cause persistent infections for 10 weeks in outbred Swiss mice [50]. K. Spindler and colleagues then showed that MAV-1 infections persist for at least 55 weeks in outbred Swiss mice [51]. In addition, Spindler and colleagues demonstrated that in contrast to humans, mice that lack B cells are highly susceptible to MAV-1 infection, while mice that lack T cells are not susceptible [52]. In light of our results, the question could be raised as to whether anti-MAV-1 antibodies are needed to generate T_regs_ to reduce the potential impact T-cell induced immunopathology [27]. To address this one could use a single preinjection of sera from MAV-1-challenged mice into B-cell deficient mice and quantify disease progression. Using nonhuman primates (NHPs) to address the dichotomy between the two arms of the adaptive immune response to adenoviruses is likely a more informative option, but use of NHPs entails unique challenges when it comes to preexisting exposure to their own set of adenoviruses. Nonetheless, Miller and colleagues elegantly showed that NHPs, harboring neutralizing antibody response against a HAdV5 host-range mutant, and then re-challenged with the same virus, respond with a significant increase in circulating T_regs_ [53]. These *in vivo* observations, which hinge on the pre-existing neutralizing antibodies, are consistent with our proposed mechanism. One also needs to take into the dynamic, recurrent exposure to HAdVs during childhood and adolescence. These encounters provide multiple opportunities for the formation of IC-HAdVs and the impetus to form HAdV-specific tolerogenic DCs and T_regs_.

Our data also complement the mechanism for HAdV5 persistence described by Hearing and colleagues [6]. Using human cell lines, they showed that IFN-α and IFN-γ production block HAdV5 replication via an E2F/Rb transcriptional repression of its E1A immediate early gene [54]. The E1A gene product is essential for activating expression of the other early genes and reprogramming the cell into a state that allows virus propagation. Of note, type 1 IFN secretion is significant from IC-HAdV5-challenged DCs and may allow HAdVs (including those that are covered with non-neutralizing Abs) to be taken up by neighboring cells to establish persistent infections.

Mechanisms by which DCs promote tolerance include induction of T_regs_, the inhibition of memory T-cell responses, T-cell anergy, and clonal deletion [24–26]. The semi-mature phenotype of tolerogenic DCs provide insufficient stimulatory signals and drive naïve T cells to differentiate into T_regs_ rather than effector T cells [55]. The global anti-viral response by DCs acts via a combinatorial cytokine code to direct the response of neighboring immune cells. The cytokine profile produced by the IC-HAdV5-challenged DCs and bystander DCs is noteworthy, particularly in the context of the combination and dose that influences activation of other immune cells. Recently, a biochemical and functional chemokine interactome study suggested that several chemokines form heterodimers that have unique functions in certain conditions [56]. Based on these interactome data, we plotted the possible combinations that could influence the direct and bystander DCs in our assays (**Figure S11**). What impact these heterodimers could have on HAdV persistence will require future study, in particular because we did not find notable levels of TGFβ secreted by direct or bystander DCs. More than other cytokine families, the IL-1 family may be primordial because it is tightly linked to IC-HAdV-induced DC pyroptosis. Indeed, the intracellular domain of the IL-1R1 shares similar signaling function properties with TLRs. In general, IL-1β release from monocytes is tightly controlled; less than 20% of the total pro-IL-1β precursor is processed and released. IL-1β also increases the expression of intercellular adhesion molecule-1 and vascular cell adhesion molecule-1, which, together with the chemokines, promote the infiltration of cells from the circulation into the extravascular space and then into inflamed tissues [57]. While circulating monocytes do not constitutively express *IL1β* mRNA, adhesion to surfaces during diapedesis induces the synthesis of large amounts that are assembled into large polyribosomes primed for translation [58].

Two aspects of the IC-HAdV-induced DC immune response that remain unknown are the impact of neutrophils and the phenotype/function of recruited monocytes. Neutrophils are pertinent because they can secrete/release proteinase 3 (PR3), elastase, cathepsin-G, chymase, chymotrypsin, and meprin α or β, which can process extracellular pro-IL-1β into its active form [59,60]. In addition, IC-HAdVs activate neutrophils (L-selectin shedding) via Fc receptors and complement receptor 1 interactions [61]. Moreover, neutrophils are a major source for anti-microbial peptides (e.g., defensins and LL-37) and proteins (e.g. lactoferrin) for which a pro- or anti-viral roles in HAdV infection has been proposed [62]. With respect to the phenotype/function of recruited monocytes, Ly6C^hi^ monocytes [63], which suppress T-cell proliferation during HAdV-induced inflammation [64], may also impact the creation of HAdV antigen-presenting tolerogenic DCs and HAdV-specific T_regs_.

The dynamic equilibrium between recurrent HAdV infections and IC-HAdV formation, DC maturation/pyroptosis, recruitment and generation of bystander DC, and T_regs_ production/activation, likely starts in childhood and develops nonlinearly over decades. While it is hard to argue that the generation of persistent infections is not beneficial to the pathogen, it is possible that the sustained anti-HAdV cellular and humoral responses partially shield a healthy host from infections by other pathogens (e.g. hepatitis C virus [65]) or the related immune-induced tissue damage [66]. Avoiding chronic tissue damage is particularly important because, as mentioned previously, HAdVs infect the eye, respiratory and gastrointestinal tracts. However, in a T-cell compromised host IC-HAdV-induced pyroptosis of FcγR^+^ cells (neutrophils, monocytes, macrophages, DCs) may also prime the host for HAdV-disseminated disease.

In summary, our findings suggest a mechanism by which humoral immunity to HAdV fosters tolerance. Understanding this complex virus-host interplay may enable us to identify high risk patients undergoing immunosuppression and develop therapies to treat disseminated HAdV-disease [67,68].

## Materials and Methods

### Ethics statement

Blood samples from >120 anonymous donors from the local blood bank (Etablissement Français du sang, Montpellier, France) were used during this study. All donors provided written informed consent.

### Cells and culture conditions

DCs were generated from freshly isolated CD14^+^ monocytes in the presence of 50 ng/ml granulocyte-macrophage colony-stimulating factor (GM-CSF) and 20 ng/ml interleukin-4 (IL-4) (PeproTech, Neuilly sur Seine, France) [3]. DC stimulations were performed 6 days post-isolation of monocytes. THP-1 cells purchased from ATCC (TIB-202) were cultured in RPMI 1640 medium supplemented with 10% fetal bovine serum (FBS). Similar to DCs, THP-1 cells were differentiated into DCs using 50 ng/ml GM-CSF and 20 ng/ml IL-4 for 6 days.

### HAdV vectors & hexon peptides

Adβgal is a ΔE1/E3 HAdV5 vector harboring a *lacZ* expression cassette [69]. Ad^L40Q^ is an HAdV5-based vector with a leucine to glutamine mutation of an amino acid in protein VI that decreases its membrane lytic activity [31]. Alexa555- and Alexa488-HAdV5 were generated from Adβgal by using an Alexa555 or Alexa488 Protein Labeling Kit (Life Technologies, Villebon-sur-Yvette, France) as previously described [70]. Ad2ts1 harbors a mutation in protease and results in several unprocessed capsid proteins and a hyper-stable capsid [71]. All HAdV viruses/vectors were produced in 293 or 911 cells and purified by double banding on CsCl density gradients as previously describe [14]. Vector purity typically reaches >99%. HAdV concentrations (physical particles/ml) were determined as previously described [72]. The hexon peptide pool (PepTivator AdV5 hexon, Miltenyi) is overlapping sequences of the HAdV5 hexon protein.

### Antibodies

Anti-human CD4-PE, anti-human CD83-FITC (cat 556910), anti-human HLA-ABC-PE (cat 555553), anti-human HLA-DR-PE (cat 555812), anti-human CD80-FITC (cat 557226), anti-human CD86-APC (cat 555660), anti-human CD25-PE (cat 555432), anti-human CD127-FITC (cat 561697), anti-human CD4-PE-Cy7 (BD) (cat 348809), anti-TNF-PE-Cy7 (cat 557647), anti-IL-2-PE (cat 554566), anti-IFN-γ-APC (cat 554702), anti-IL-10-PE (cat 554706) were from Becton Dickinson, Pharmigen. Anti-human Foxp3-APC (cat 17-4776-41) was from eBioscience. Anti-human CD14-PE (cat A07764) was from Beckman Coulter. Anti-human CD45RO-APC/Cy7 (cat 304227), anti-human CD45RA-PE (cat 304205), anti-human CD3-APC (cat 300411), and anti-human CD40-APC (cat 313008) were from BioLegend).

### Immune complex formation and DC stimulations

DCs (4 × 10^5^ in 400 µl of complete medium) were incubated with HAdV5 or IC-HAdV5 (or IC) (2 × 10^4^ physical particles (pp)/cell, unless otherwise indicated) for the indicated times. IC-HAdV5s were generated by mixing the virus (8 × 10^9^ physical particles) with 2.5 µl of IVIg (human IgG pooled from 1,000 to 50,000 donors/batch) (Baxter SAS, Guyancourt, France) for 15 min at room temperature. IVIg is used in patients with primary or acquired immunodeficiency as well as autoimmune diseases. Z-VAD-FMK 20 µM (ZVAD) was added 2 h before stimulation. Brefeldin A was used at 3 µg/ml after 6 h stimulation or for the same time with stimulation.

### Bystander DC stimulation

DCs (1.5 × 10^6^ in 1.5 ml of full media) were incubated ± LPS 100 ng/ml, HAdV5, and IgG in the lower compartment of the well (12 mm diameter polyester membranes with 0.4 µm pores; (Corning, Bagneaux-sur-Loing, France). After 6 h incubation, fresh immature DCs (6 × 10^5^ in 600 µl of media) were added to the upper compartment and are referred to as bystander DCs. TAK-242 was added to DCs 1 h pre-challenge.

### Quantification of mRNA

Expression levels of cytokine and chemokine genes were evaluated using RT-qPCR assays. Total RNA was isolated from cells using the high pure RNA isolation Kit (Roche, Berlin, Germany) with a DNase I treatment during the purification and subsequent elution in 50 µl of RNase-free water (Qiagen, IN, USA). Reverse transcription was performed with the superscript first-strand synthesis system (Invitrogen) using 10 µl of total RNA and random hexamers. The cDNA samples were diluted 1:20 in water and analyzed in triplicate using a LightCycler 480 (Roche, Meylan, France). SYBR green PCR conditions were as follows: 95°C for 5 min and 45 cycles of 95°C for 15 s, 65°C or 70°C for 15 s, and 72°C for 15 s using *GAPDH* as a standard. See **Table S2** for primers sequencers. Relative gene expression levels of each respective gene were calculated using the threshold cycle (2^-ΔΔCT^) method and normalized to *GAPDH* mRNA levels.

### RT^2^ Profiler ^TM^ PCR array

Expression levels of cytokine and chemokine mRNAs were analyzed using PCR array assays. Total RNA was isolated from cells using the High Pure RNA isolation Kit (Roche, Berlin, Germany) with a DNase I treatment during the purification and elution in 50 µl of RNase-free water (Qiagen). Reverse transcription was performed with the RT^2^ First strand Kit (Qiagen, Courtaboeuf, France), and the cDNA samples were analyzed in duplicate using a RT^2^ Profiler ^TM^ PCR array (Qiagen). SYBR green PCR conditions were 95°C for 10 min and 40 cycles of 95°C for 15 s, and 60°C for 1 min using 84 human inflammatory and receptor genes. The potential mRNAs were chosen and then confirmed by RT-qPCR.

The genes that contributed in each axis in the PCA were as follows: **F1** = *CCL1, 2, 4, 5, 7 13, 15, 17, 20, 22, CSF1, CX3CL1, CXCL 1 to 3, 5, 8 to 11, FASLG, IFNG, IL10RA, IL10RB, IL15, IL1a, IL1b, IL7, NAMPT, TNFSF4, 10, 11, 13, 13B*, and *VEGFA*. **F2** = *AIMP1, C5, CCL1, 2, 13, 17, 23, CRR1, 2, 3, 4, 5, CSF1, CX3CR1, CXCR2, IL10RA, I10RB, IL15, LTA, LTB, MIF, SPP1, TNF, TNFSF4, 10, 11, 13*, and *13B*. **F3** = *CCL17, 23, CCR5, CX3CR1, IL10RA, IL5, IL9, MIF*, and *OSM*.

### Co-stimulatory protein levels

Surface levels of CD83, MHCII, CD80, CD40, and CD86 were assessed by flow cytometry. Cell membrane integrity was assessed by collecting cells via centrifugation at 800x g; the cell pellets were then resuspended in PBS containing 10% FBS, propidium iodide (PI) (Sigma-Aldrich, Missouri, USA), or 7-aminoactinomycin D (7AAD) (Becton-Dickinson, New Jersey, USA). The cell suspension was incubated for the indicated times and analyzed using a FacsCalibur flow cytometer (Becton-Dickinson) and FlowJo software.

### Intracellular staining

Surface and intracellular levels of CD83 and CD86 (total protein) were stained with a BD Cytofix/Cytoperm™ Fixation/Permeabilization Kit, and then measured by flow cytometry. To assess cell membrane integrity, the cells were collected and centrifuged at a speed of 800x g; the cell pellets were then resuspended in PBS, 10% FBS, PI (Sigma), or 7AAD and analyzed on a FacsCalibur flow cytometer (Becton-Dickinson) and FlowJo software.

### Monocyte migration assay

Monocyte migration was evaluated using a 5.0 µm-diameter pore transwell system (Corning, Bagneaux-sur-Loing, France). Monocytes (2 × 10^5^ in 200 µl of full media) were added into inserts and DCs or DCs (7.5 × 10^5^ in 750 µl of full media) and ± LPS (100 ng/ml), HAdV5, HAd555, or IgG in the lower wells. Monocytes were stained by carboxy-fluorescein diacetate-succinimidyl ester (CFSE) (Molecular Probes, Eugene, OR, USA) (CellTrace™ CFSE Cell Proliferation Kit). DCs incubated for 30 min with HAd555 or HAdV5 and IgG in the lower chamber were or were not washed in medium before adding the stained CFSE monocytes. After 3, 6, and 24 h incubation at 37°C, the cells in the upper and lower compartment were detected quantified using a FacsCalibur flow cytometer (Becton-Dickinson) and FlowJo software.

### Cytokine secretion: ELISA and Luminex assays

Supernatant from the cells were collected and cytokine secretion was measured by ELISA and Luminex assays. The secretion of TNF and IL-1β was quantified by ELISA using an OptEIA human TNF ELISA Kit (Becton Dickinson) and human IL-1β/IL-1F2 DuoSet ELISA (R&D Systems, Lille, France) following the manufacturer’s instructions. Additionally, 22 other cytokines and chemokines were detected by Luminex using a Bio-plex pro human chemokine, cytokine kit (Bio-Rad, Marnes-La-Coquette, France) following the manufacturer’ instructions.

### Depletion of CD25^+^ from PBMC

PBMC were isolated using standard gradient separation techniques. Half were CD25^+^-depleted, using anti-CD25 in a human CD4^+^CD25^+^CD127^dim/-^ Regulatory T Cell Isolation Kit II and MACS separation system.

### CFSE and VPD 450 labeling

PBMCs were washed and suspended in PBS for labeling with CFSE or Violet Proliferation Dye 450 (VPD 450) (BD Horizon™, Le Pont de Claix, France) at a final concentration of 2.5 μM or 1 μM, respectively, for 3 min at room temperature. Labeling was terminated by the addition of fetal calf serum (FCS) (40% of total volume).

### PBMC activation assays

PBMCs ± CD25^+^ were stained with CFSE and cultivated in 96-well U-bottom plates; (cell concentration 1 × 10^6^/ml and a final volume of 200 μl; PBMC CD25^+^/ PBMC CD25^-^ ratio 1:10). HAdV5 hexon peptides (PepTivator, Miltenyi, Paris, France) were added at 0.3 nmol. On days 3 and 5 the cells were split and IL-2 was added (final concentration 100 U/ml). Cells were analyzed on a FACS Canto II using FlowJo software.

### T_reg_ generation

Naïve CD4^+^ T cell were isolated using naïve CD4^+^ T Cell Isolation Kit II and MACS separation system. DCs indirectly activated for 12 h with LPS, HAdV5, IC-HAdV5 and IgG, and then were co-cultured with CD4^+^ naïve T cells labeled VPD450 (with ratio bystander DCs/ T cells is 3:1) in RPMI 1640 supplemented with 10% FCS and IL-2 (Proleukin 18 × 10^6^ IU, CA, USA) (100 U/ml) for 3 or 7 days. Recombinant IL-2 was added on day 3 and day 5. CD25, CD127, and FoxP3 levels were quantified by flow cytometry using FACS Canto II.

### Statistical analyses

All experiments were performed at least in duplicate a minimum of three independent times, and the results are expressed as mean ± SEM unless otherwise stated. The statistical analyses were performed using the Student’s *t*-test unless otherwise stated. A *p* value < 0.05 is denoted as significant. Statistical analyses of the global cytokine profiles (pie chart) were performed by partial permutation tests using the SPICE software.

## Data availability

All data generated or analyzed during this study are included in this published article (and its supplementary information files).

## Acknowledgments

We thank Sylvie Grandemange, Fabien Blanchet, Sebastian Nisole, Valerie Dardalhon, and Claire Daien for reagents and advice. We thank EKL members for technical help and constructive comments. We thank the MRI, member of the national infrastructure France-BioImaging, and SERENAD for statistical analyses. KE current address: Vaccine and Infectious Disease Division, Fred Hutchinson Cancer Research Center, Seattle, WA, USA.

## Supporting information

**Figure S1).**
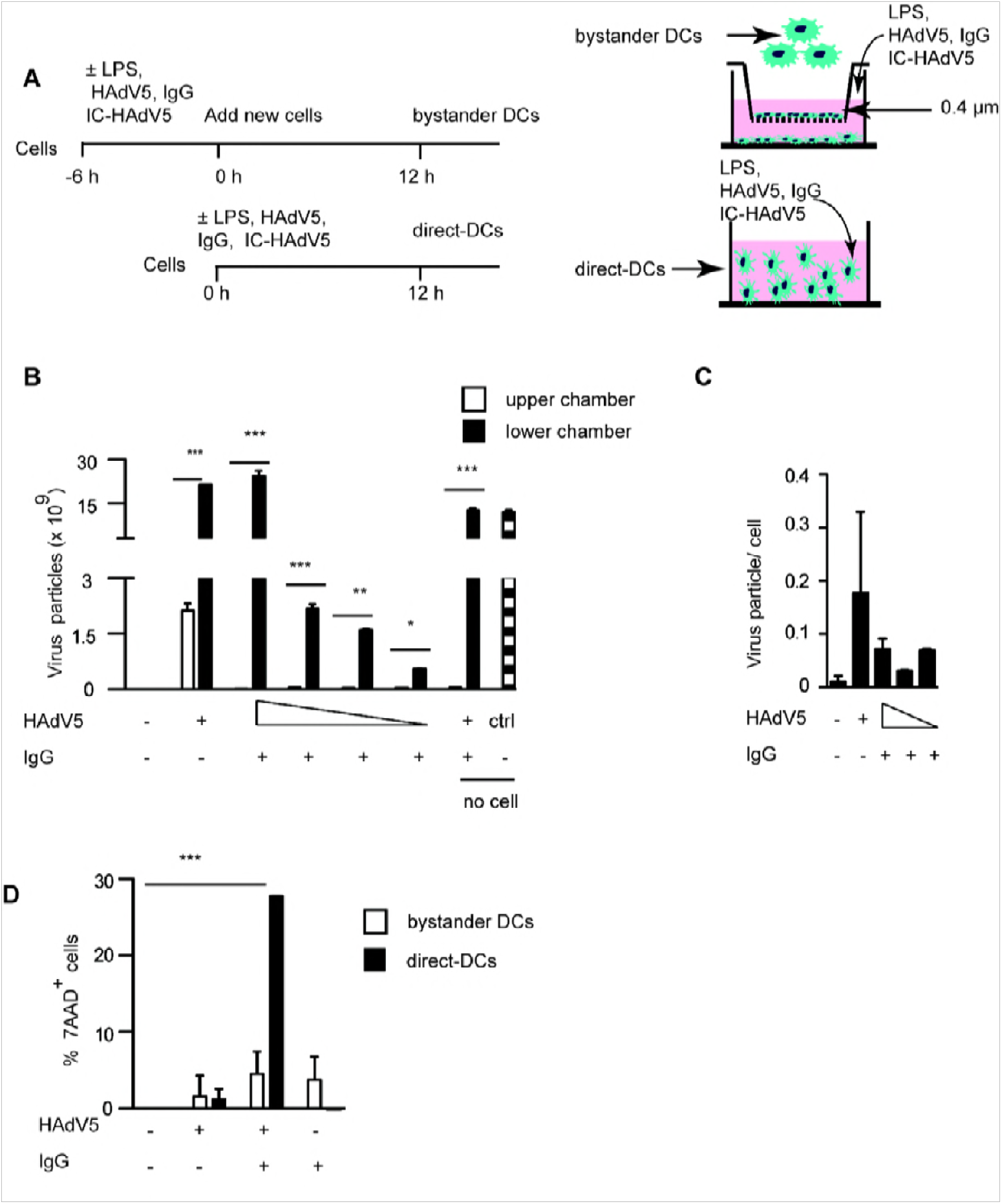
Transwell assay setup and controls. **A)** We used transwell inserts with 0.4 um filter to generate direct and bystander DCs. Direct DCs (1.5 ×10^6^ cells unless mentioned otherwise) were incubated with the stimulus (e.g. LPS, HAdV5, mutant virus, ± IVIg ± drugs) in lower compartment for 6 h. Fresh DCs (6 × 10^5^) were added to the upper compartment. **B)** To determine if HAdV5 particles (2 × 10^4^ pp/ml) added to the lower chamber diffused to the upper compartment and impact the bystander DCs, we quantified (by qPCR) HAdV5 genomes in the **supernatant** of each compartment. 1.6 × 10^10^ pp of HAdV5 pp were used in the control medium. These data demonstrate that 10,000-fold fewer particles could be found in the upper chamber. **C)** Quantification of HAdV5 genomes **associated with bystander DCs** as measured by qPCR (n ≥3). DNA from mock-treated samples was extracted and virus/cell was normalized to *GAPDH* copy number. The quantity of HAdV5 genomes/cell was normalized by *lacZ* (transgene in the vector) vs. *GAPDH* copy number. While direct DCs take up ∼600 pp/cell [14], we found that 1 in 10 bystander DC contains a single HAdV5 genome. **D)** The 7AAD^+^ bystander and direct DCs (i.e. DCs with compromised plasma membrane integrity) in each condition were quantified by flow cytometry. The assays were carried out in 4 donors (mean ± SEM. These results demonstrate that bystander DCs do not show loss of cell membrane integrity. *p* values were derived using Student’s t-test (**B & C**) or one-way ANOVA with Dunnett’s post-tests (**D**). * *p <* 0.05, ** *p <* 0.01 and *** *p <* 0.001.

**Figure S2).**
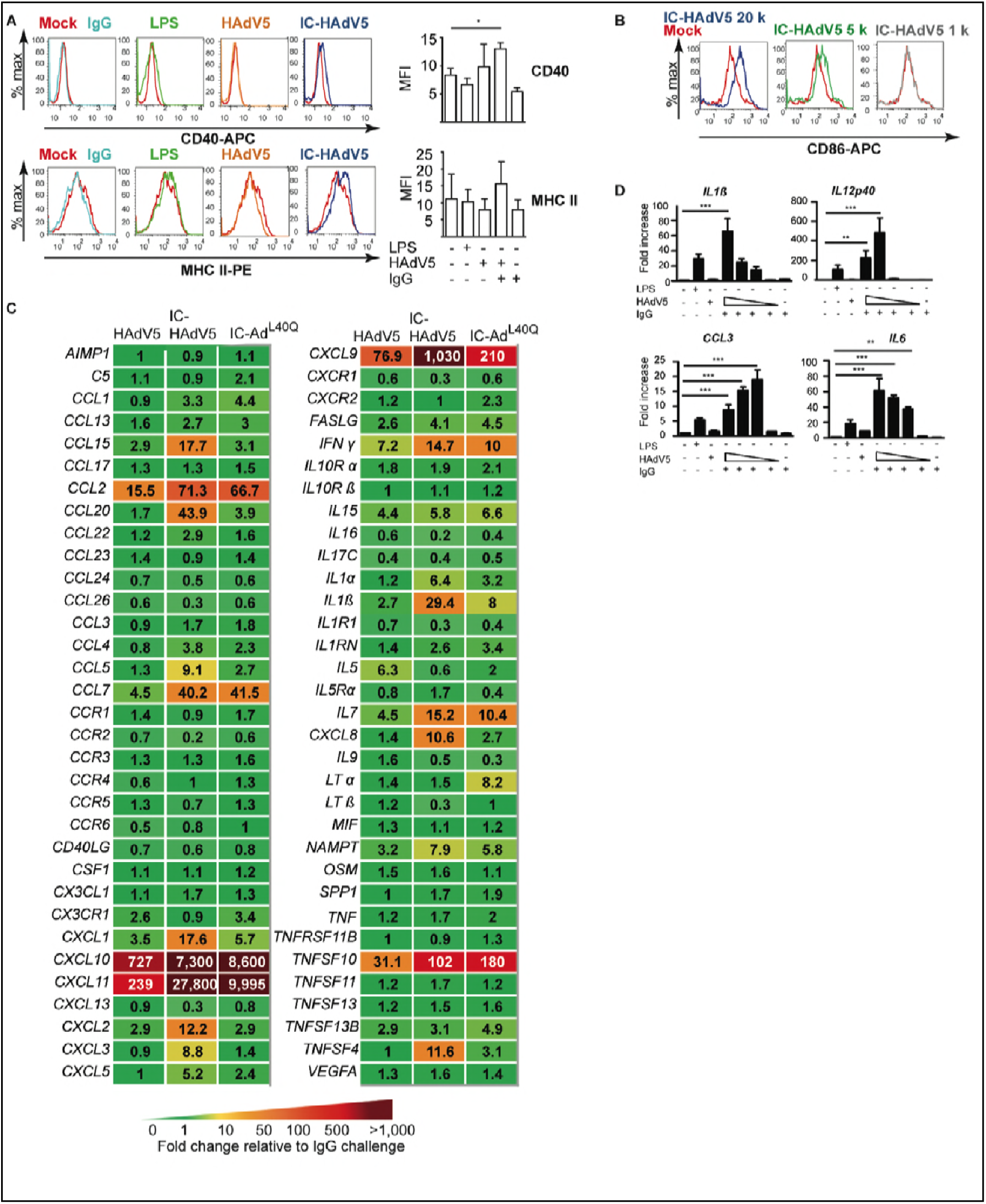
Maturation/activation markers on bystander DCs. Bystander DCs were generated using milieu from DCs challenged with IgG, LPS, HAdV5, or IC-HAdV5. The color code is as in **Figure 2**. **A)** The data are representative flow cytometry profiles of CD40 and MHC II surface expression. A modest increase was noted in each case. **B)** In a dose-dependent assay (20,000, 5,000, or 1,000 pp/cell) CD86 cell surface levels were quantified detected by flow cytometry. The data are representative flow cytometry profiles. Assays were carried out in 4 donors with similar results. **C)** PCR array profiles from bystander DCs exposed to the milieu generated by DCs challenged by HAdV5, IC-HAdV5, and IC-Ad^L40Q^. The 66 cytokine mRNAs that gave unique qPCR peaks in our hands. **D)** *IL1β, IL12p40, CLL3* and *IL6* mRNA levels in bystander THP1 DCs assayed in a dose-dependent (20,000, 10,000, 5,000, or 1,000 pp/direct DC) response. Data are mean ± SEM with 3 independent experiments. *p* values were derived from one-way ANOVA with Dunnett’s test. * *p <* 0.05, ** *p <* 0.01 and *** *p <* 0.001.

**Figure S3).**
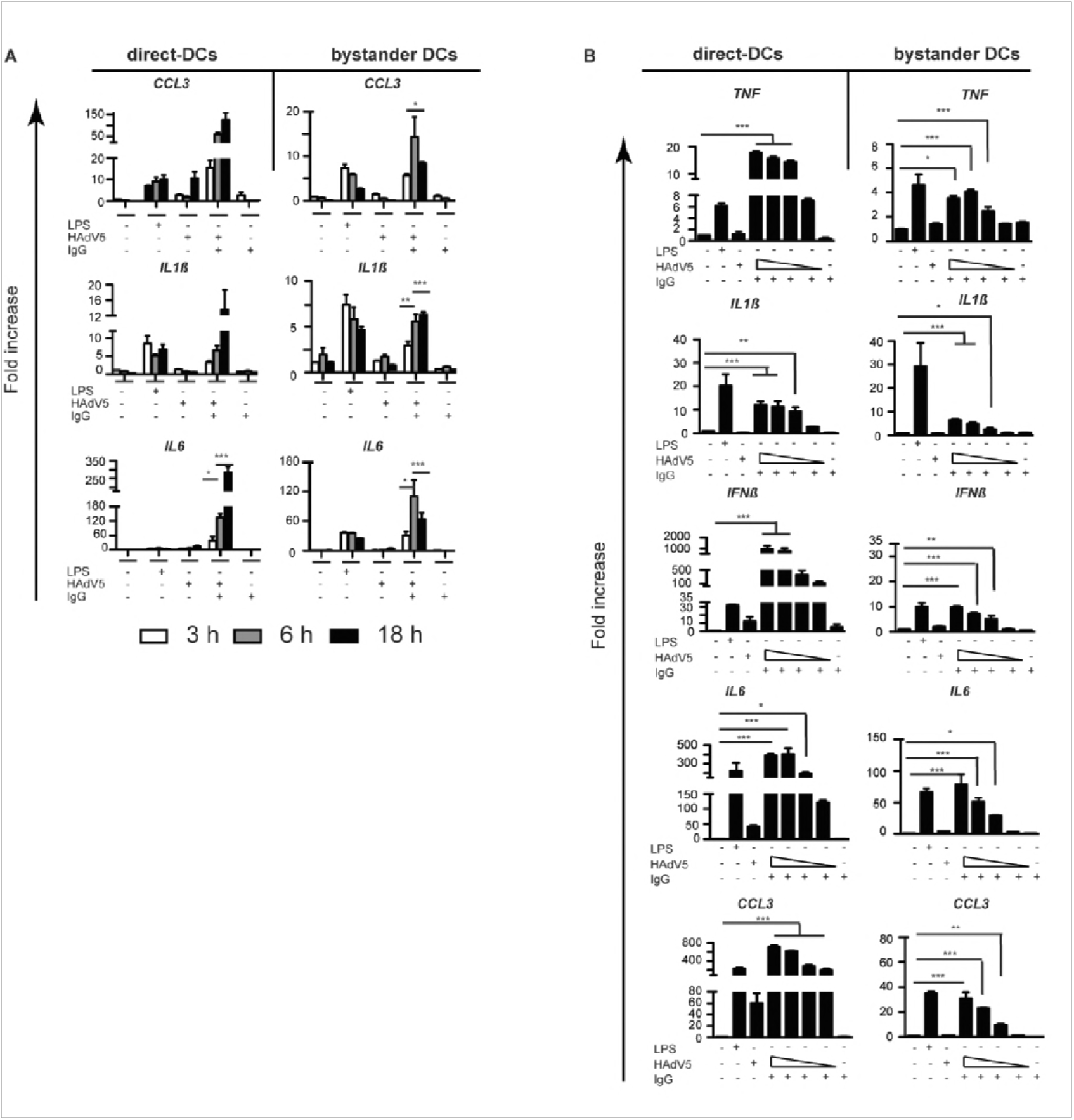
Bystander and direct DC cytokine mRNA levels as a function of time and dose. We extended the mRNA array results by quantifying dose-dependent responses of a handful of mRNA levels by RT-qPCR. Because DCs derived from monocytes from random blood bank donors can have widely different levels of mRNAs, we compared mRNA levels in THP-1-derived DCs to provide a standardized view of the changes. THP-1 cells were differentiated into DCs for 6 days, then directly and indirectly activated. **A)** *CCL3, IL1β*, and *IL6* mRNA levels in DCs challenged with LPS, IgG, HAdV5 and IC-HAdV5 (left hand column), and bystander DCs (right hand column) incubated in the respective direct DC milieu were quantified at 3, 6, and 18 h post-incubation. **B)** Changes in *TNF, IL1β*, IFN*β, IL6, and CCL3* mRNA levels in direct (left hand column) *TNF*: IC 2 × 10^4^ vs. 10^4^ ns; 10^4^ vs. 5 × 10^3^ ns; 5 × 10^3^ vs. × 10^3^ ***; *IL1β* IC 2 × 10^4^ vs. 10^4^ ns; 10^4^ vs. 5 × 10^3^ ns; 5 × 10^3^ vs. 10^3^ ns; IC 2 × 10^4^ vs. 10^3^ **, IC 10^4^ vs. 10^3^ *; *IFNβ:* IC 2 × 10^4^ vs. 10^4^ ns; 10^4^ vs. 5 × 10^3^ ns; 5 × 10^3^ vs. 10^3^ ns, IC 2 × 10^4^ vs. 10^3^ *; *IL6*: IC 2 × 10^4^ vs. 10^4^ ns; 10^4^ vs. 5 × 10^3^ **; 5 × 10^3^ vs. 1 × 10^3^ ns; *CCL3*: IC 2 × 10^4^ vs. 1 × 10^4^ ns; 1 × 10^4^ vs. 5 × 10^3^ ***; 5 × 10^3^ vs. 10^3^ ns) Bystander DC (right hand column) dose-dependent assay (2 × 10^4^, 10^4^, 5 × 10^3^, or 10^3^ pp/cell) by RT-qPCR *TNF*: IC 2 × 10^4^ vs. 10^4^ ns; 10^4^ vs. 5 × 10^3^ ns; 5 × 10^3^ vs. 1 k ns, IC 2 × 10^4^ vs. 10^3^ **, IC 10^4^ vs. 10^3^ ***; *IL1β:* IC20 k vs. 10^4^ ns; 10^4^ vs. 5 × 10^3^ ns; 5 × 10^3^ vs. 10^3^ ns; *IFNβ:* IC 20 k vs. 10^4^ ns; 10^4^ vs. 5 × 10^3^ ns; 5 × 10^3^ vs. 10^3^ **; *IL6:* IC 2 × 10^4^ vs. 10^4^ ns; 10^4^ vs. 5 × 10^3^ ns; 5 × 10^3^ vs. 10^3^ ns, 10^4^ vs. 5 × 10^3^ *** *CCL3*: IC 2 × 10^4^ vs. 10^4^ ns; 10^4^ vs. 5 × 10^3^ ***; 5 × 10^3^ vs. 10^3^ *). As in “**A**” controls included IgG and HAdV5. Three independent experiments were carried out. Data are mean ± SEM. *p* values were derived using Student’s *t*-tests. * *p <* 0.05, ** *p <* 0.01 and *** *p <* 0.001..

**Figure S4).**
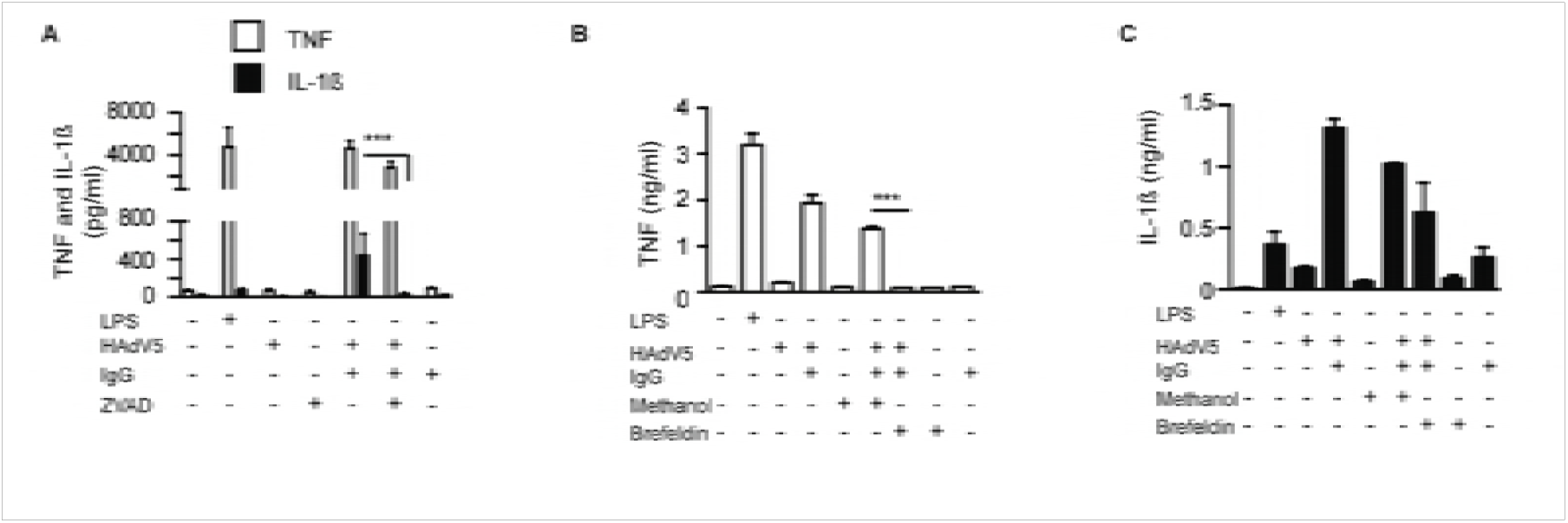
Controls for ZVAD and brefeldin A assays. **A)** TNF and IL-1β secretion in response to ZVAD treatment (2 h before challenge) of DCs challenged with LPS, IgG, HAdV5, and IC-HAdV5. **B)** DCs were simultaneously treated with brefeldin A and challenged with LPS, IgG, HAdV5, and IC-HAdV5. TNF secretion was quantified at 18 h. **C)** DCs were simultaneously treated with brefeldin A and challenged with LPS, IgG, HAdV5, and IC-HAdV5. IL-1β secretion was quantified at 18 h. Data are mean ± SEM, *p* values were derived from Student’s *t*-tests, n ≥ 3 donors. *** *p <* 0.001.

**Figure S5).**
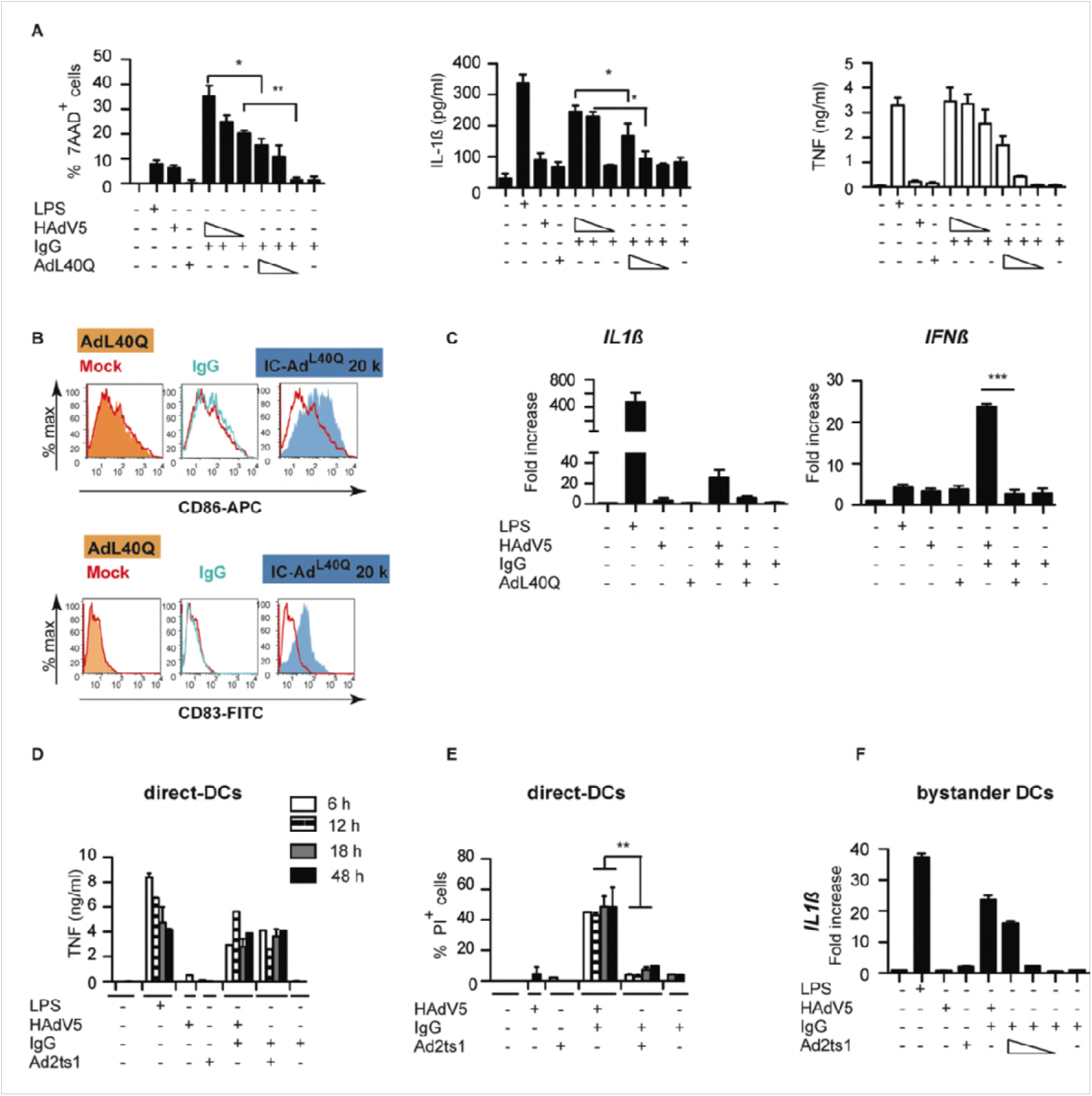
Controls for IC-Ad^L40Q^ and IC-Ad2ts1. **A)** DCs challenged with LPS, IgG, HAdV5, Ad^L40Q^ and increasing concentrations of IC-HAdV5 and IC-Ad^L40Q^ were analyzed for loss of membrane integrity (7AAD^+^ cells), IL-1β and TNF secretion. **B)** Cell surface levels of the maturation/activation markers CD86 and CD83 following direct DCs challenged with IgG, Ad^L40Q^, IC-Ad^L40Q^, HAdV5, and IC-HAdV5. **C)** bystander DC *IL1β* and *IFNβ* mRNA levels quantified by RT-qPCR assay. Experiments were carried out in ≥3 donors. *p* values were derived from Student’s *t*-tests. *, **, *** denote *p* values of < 0.05, < 0.01, < 0.001, respectively. DCs were challenged with LPS, IgG, HAdV5, IC-HAdV5, Ad2ts1, and IC-Ad2ts1 and screened for **D)** time-dependent (6 to 48 h) TNF secretion; and **E)** time-dependent (6 to 48 h) loss of membrane integrity using propidium iodide (PI) incorporation; or **F)** DCs were challenged with LPS, IgG, HAdV5, IC-HAdV5, Ad2ts1, and IC-Ad2ts1 and then used to generate bystander DCs in which the *IL1β* mRNA levels were quantified by RT-qPCR assay following dose-dependent stimulation (20 × 10^3^, 10 × 10^3^, or 5 × 10^3^ pp/cell) of the direct DCs. All experiments were carried out in 3 donors and in duplicate. *P* values were derived from Student’s *t*-tests. ** *p <* 0.01.

**Figure S6).**
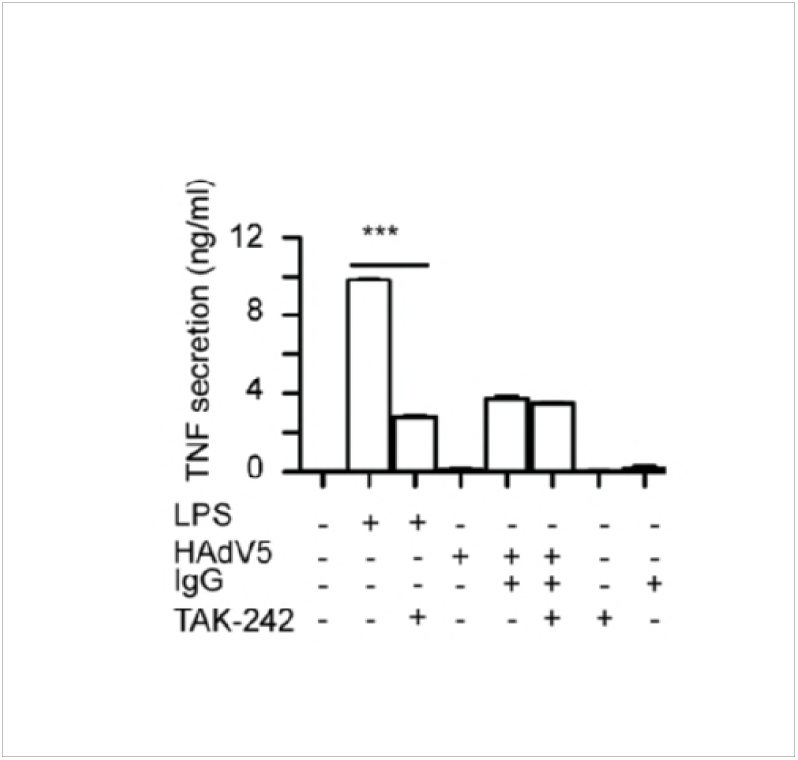
TAK-242 controls. Bystander DCs were treated with TAK-242 for 1 h before adding them to the DCs challenged with LPS, IgG, HAdV5, or IC-HAdV5. TNF secretion was quantified in direct DCs in the lower compartment (n = 3 donors). *p* values were derived from Student’s *t*-tests. *** *p <* 0.0001.

**Figure S7).**
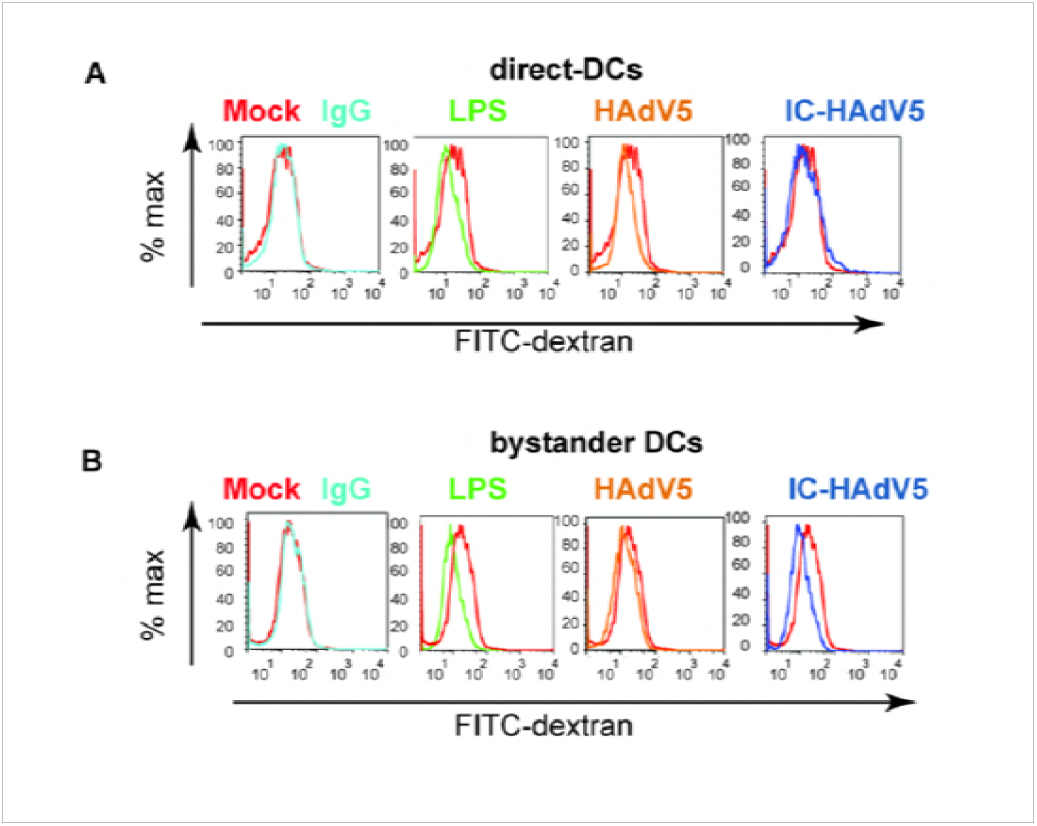
Controls for fluid phase uptake assay. Nonspecific binding of dextran to **A)** direct DCs and **B)** bystander DCs was controlled by incubating DC (post-stimulation) with FITC-labeled dextran at 4°C. Direct DCs were challenged with IgG, LPS, HAdV5, or IC-HAdV5. The cells were then incubated with FITC-labeled dextran and analyzed by flow cytometry. The data are representative flow cytometry profiles with experiments performed using cells from in 3 donors and in duplicate.

**Figure S8).**
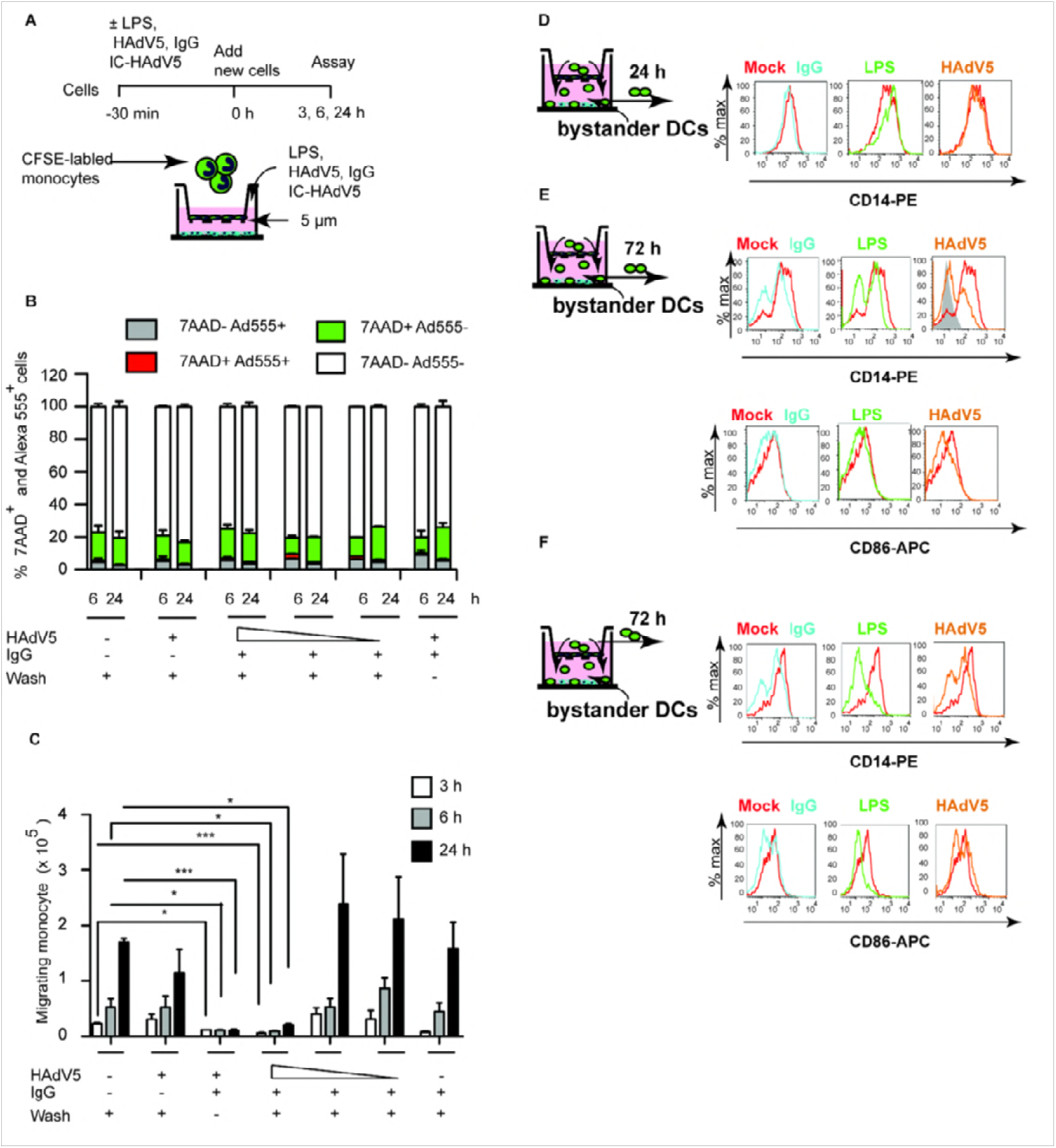
Controls for monocyte recruitment. **A)** A 5-micron-pore membrane transwell system was used for monocyte migration assays. The timing and stimuli are indicated in the schematics. Round green cells are CFSE-labeled monocytes. **B)** These data shown percentage of monocyte in the upper chamber that potentially interact with HAdV or IC-HAdV5. **C)** To address this possibility, we covalently linked Alex555 to the HAdV5 capsid (HAdV5-Alexa555 [29]) to identify cells associated with HAdV5 or IC-HAdV5. CFSE-labeled monocytes were then assayed by flow cytometry for loss of membrane integrity (7AAD^+^ cells) and the presence of HAdV5-Alexa555 at 6 and 24 h. These data demonstrate that ICs do not go through the pore to interact with monocytes in the upper chamber. **D)** CD14 expression levels on monocytes recruited towards bystander DCs that were created with the milieu from DCs challenged with IgG, LPS or HAdV5 at 24 h. **E)** CD14 and CD86 levels on monocytes recruited to bystander DCs that were created with the milieu from DCs challenged with IgG, LPS or HAdV5 at 72 h. **F)** CD14 and CD86 expression levels on monocytes that remained in the upper compartment at 72 h. The lower compartment contained bystander DCs that were created with the milieu from DCs challenged with IgG, LPS or HAdV5. The data are representative flow cytometry profiles with assays carried out in 4 donors. * *p <* 0.05, ** *p <* 0.01 and *** *p <* 0.001.

**Figure S9).**
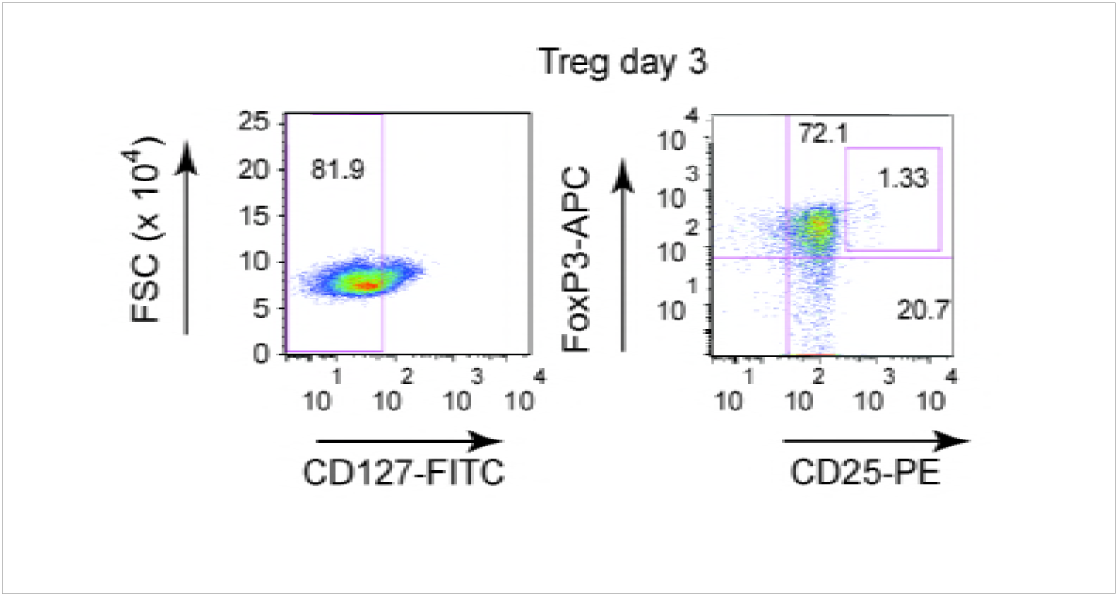
Bystander DCs, generated using the media from DCs challenged with IgG, HAdV5, and IC-HAdV5, were incubated with naive CD4^+^ T cells isolated from the same donors. Three days post-incubation we gated on CD127^dim^ cells to identify CD25^+^/FoxP3^high^ cells. The data are representative flow cytometry profiles with assays carried out in 7 donors.

**Figure S10).**
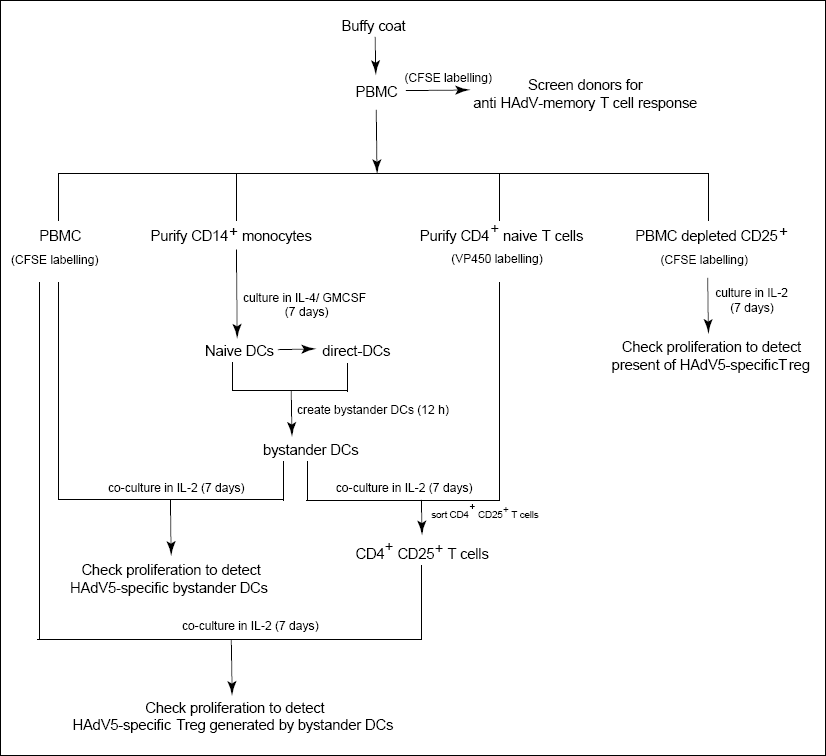
Flow chart demonstrating the cells and process used for the T_reg_ assays

**Figure S11).**
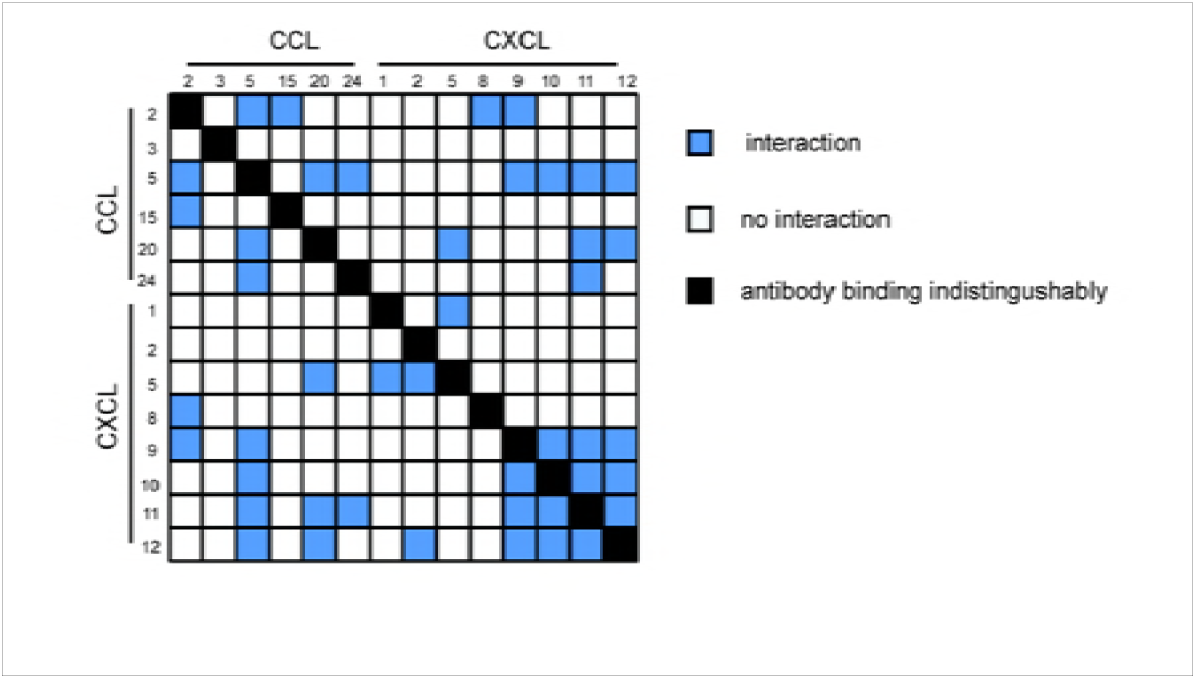
Potential cytokine heterodimer formation/interactions. Potential cytokine heterodimers are based on von Hundelshausen *et al.* [56] interactome data and the response generated by direct and bystander DCs.

**Table S1).**
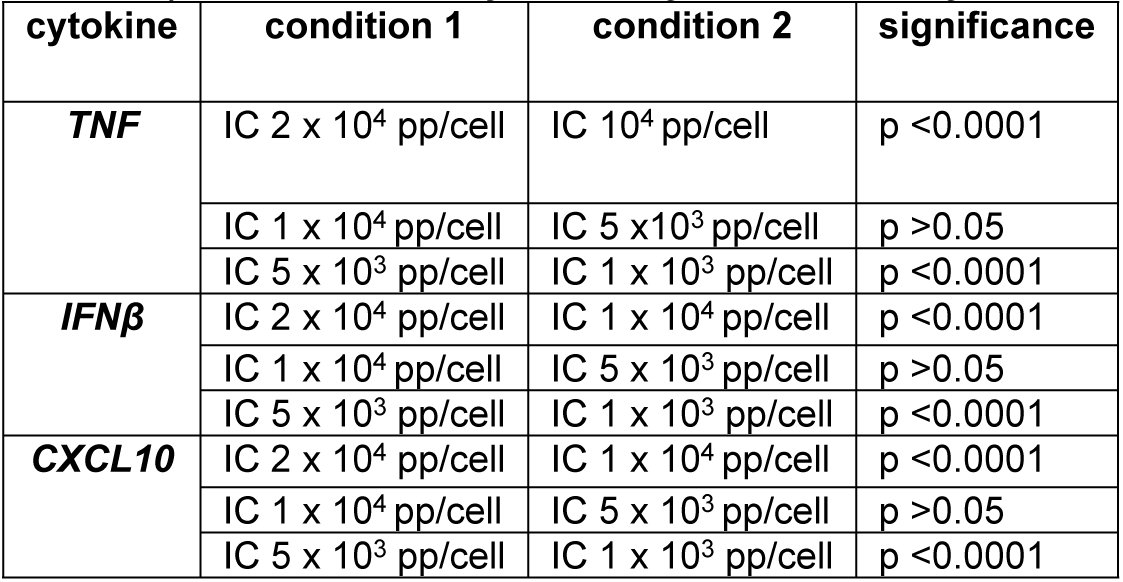
Statistical analyses of bystander DC cytokine transcription profile.

| cytokine | condition 1 | condition 2 | significance |
| --- | --- | --- | --- |
| *TNF* | IC 2 x 10<sup>4</sup> pp/cell | IC 10<sup>4</sup> pp/cell | p <0.0001 |
|  | IC 1 x 10<sup>4</sup> pp/cell | IC 5 x10<sup>3</sup> pp/cell | p >0.05 |
|  | IC 5 x 10<sup>3</sup> pp/cell | IC 1 x 10<sup>3</sup> pp/cell | p <0.0001 |
| *IFNβ* | IC 2 x 10<sup>4</sup> pp/cell | IC 1 x 10<sup>4</sup> pp/cell | p <0.0001 |
|  | IC 1 x 10<sup>4</sup> pp/cell | IC 5 x 10<sup>3</sup> pp/cell | p >0.05 |
|  | IC 5 x 10<sup>3</sup> pp/cell | IC 1 x 10<sup>3</sup> pp/cell | p <0.0001 |
| *CXCL10* | IC 2 x 10<sup>4</sup> pp/cell | IC 1 x 10<sup>4</sup> pp/cell | p <0.0001 |
|  | IC 1 x 10<sup>4</sup> pp/cell | IC 5 x 10<sup>3</sup> pp/cell | p >0.05 |
|  | IC 5 x 10<sup>3</sup> pp/cell | IC 1 x 10<sup>3</sup> pp/cell | p <0.0001 |

**Table S2:**
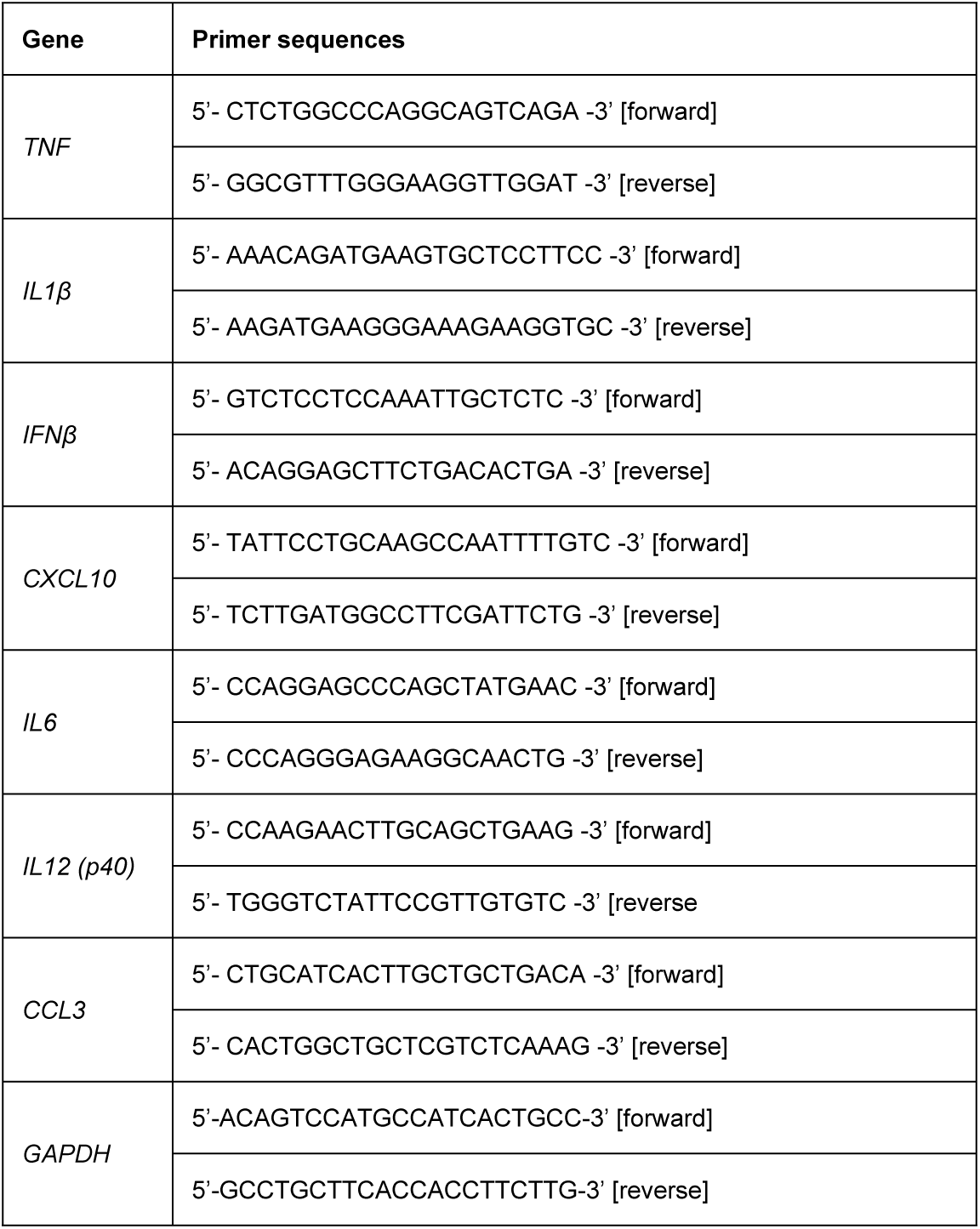
In-house designed primer sequences.

